# Significantly reduced inflammatory foreign-body-response to neuroimplants and improved recording performance in young compared to adult rats

**DOI:** 10.1101/2022.12.17.520866

**Authors:** Aviv Sharon, Maciej M. Jankowski, Nava Shmoel, Hadas Erez, Micha E. Spira

**Affiliations:** Department of Neurobiology, the Alexander Silberman Institute of Life Science, the Hebrew University of Jerusalem, Jerusalem, Israel; The Charles E. Smith Family and Prof. Joel Elkes Laboratory for Collaborative Research in Psychobiology, the Hebrew University of Jerusalem, Jerusalem, Israel; Edmond and Lily Safra Center for Brain Sciences, the Hebrew University of Jerusalem, Jerusalem, Israel

**Author notes:** Corresponding author: Micha E. Spira.

**Keywords:** Neuro-implant interface, Microglia, foreign-body-response, CSF1R-antagonists, PLX5622, PLX3397

## Abstract

The multicellular inflammatory encapsulation of implanted intracortical multielectrode arrays (MEA) is associated with severe deterioration of their field potentials’ (FP) recording performance, which thus limits the use of brain implants in basic research and clinical applications. Therefore, extensive efforts have been made to identify the conditions in which the inflammatory foreign body response (FBR) is alleviated, or to develop methods to mitigate the formation of the inflammatory barrier. Here, for the first time, we show that (1) in young rats (74±8 gr, 4 weeks old at the onset of the experiments), cortical tissue recovery following MEA implantation proceeds with ameliorated inflammatory scar as compared to adult rats (242 ±18 gr, 9 weeks old at the experimental onset); (2) in contrast to adult rats in which the Colony Stimulating factor 1 Receptor (CSF1R) antagonist chow eliminated ~95% of the cortical microglia but not microglia adhering to the implant surfaces, in young rats the microglia adhering to the implant were eliminated along with the parenchymal microglia population. The removal of microglia adhering to the implant surfaces was correlated with improved recording performance by in-house fabricated Perforated Polyimide MEA Platforms (PPMP). These results support the hypothesis that microglia adhering to the surface of the electrodes, rather than the multicellular inflammatory scar, is the major underlying mechanism that deteriorates implant recording performance, and that young rats provide an advantageous model to study months-long, multisite electrophysiology in freely behaving rats.

## 1. Introduction

Despite the overwhelming progress in neuronal engineering of implanted multielectrode arrays (MEA) [1–8], their use is impeded by the deterioration of their recording performance over time, which is manifested in a low signal-to-noise ratio (SNR), poor source resolution, as well as a drop in the number of functional implanted platforms and live electrodes [9–11]. Aside from mechanical breakdown issues that will not be discussed here [12, 13], brain implants initiate and perpetuate cascades of local noninfectious inflammatory responses culminating in the formation of encapsulating scar tissue around the implant (classically known as the ‘foreign body response’, or FBR). Based on immunohistological studies, the prevailing view holds that the FBR is the major hurdle to the effective electrophysiological use of neuroprobes over time. This view has motivated and guided diverse efforts to better understand the cell biological mechanisms underlying the formation of the FBR and research attempts to develop approaches to mitigate it.

The initiation and perpetuation of inflammatory cascades around neuroimplants are related to abiotic and biotic factors. The abiotic factors include the material from which the probes are made, their flexibility, the platform’s footprint, and its microarchitecture [14]. The biotic factors are triggered as soon as the implant is inserted into the brain tissue. Within minutes of MEA implantation, microglia residing at a distance of ~100 μm from the implant’s surface sense the event and migrate towards the implant. Three days post-implantation, activated microglia accumulate around the probes, and secrete pro-inflammatory cytokines, chemokines and reactive oxygen intermediates, which lead to neurodegenerative processes [10, 15–18]. Activated astrocytes are recruited to the encapsulating inflammatory scar by molecular signaling released from the microglia, but are delayed with respect to the microglia by approximately one week [19, 20]. The severity of the FBR depends on the specific brain site and neuron layer to which the probe was inserted [14].

Based on immunohistological imaging and electrophysiological recordings, the prevailing view attributes the recording performance deterioration of the MEA implants to three parameters. The first is the displacement of neurons away from the implants by the multicellular scar tissue [10, 19]. Because the amplitudes of the FP decline in the extracellular space at a rate of 1/r^x^, where r is the distance from the current source and x ranges from 1≤x≤2 close to the soma and x ≥2 farther from it, the amplitudes of FPs generated by displaced neurons become lower [21, 22]. The second parameter has to do with the electrical insulation of the electrodes from the displaced neurons caused by the relatively large resistivity of the multicellular scar formed around the implant which impedes the current flow towards the implant [19, 23–29]. The third parameter is the existence of reduced excitability and synaptic connectivity [30–32], demyelination [33, 34] and the degeneration of neurons in the vicinity of the implant. Attempts to experimentally verify causal relationships between the dimensions and cell density of the FBR and recording performance have not been successful. This has prompted a growing number of investigators to raise concerns as to the relevance of this dominant theory [10, 11, 20, 35–41]. Notably, a recent ultrastructural examination of the parenchyma in contact with implants revealed extensive regeneration of the neurons, including axon myelination and the formation of synaptic contacts [42].

As an alternative to the prevailing hypothesis we proposed that implant recording performance deteriorates as a result of electrode insulation caused by the adhesion of individual microglia to the electrodes’ surfaces, rather than across the high resistance multicellular encapsulating scar around the implanted MEA [20, 41–43]. One key reason why the presence of these insulating cells has been overlooked in earlier studies is that the adhering microglia were removed along with the extracted silicone-based platforms before processing the tissues for immunohistological examination. In addition, standard immunohistological analyses of the FBR traditionally use low magnification objectives which can cover the full dimensionality of the inflammatory scar tissue. Based on our observations, in which polyimide MEA platforms were thin-sectioned along with the intact brain parenchyma, and then subjected to high resolution confocal microscope imaging complemented by transmission electron microscope analysis [41, 42], we proposed that microglia adhering to the implant surfaces form a high seal resistance over the electrodes and that this seal is the underlying mechanism that impedes or blocks the electrical coupling between the neurons and the electrodes [20, 41–43]. Biophysically, this insulating mechanism is similar to the formation of a high resistance biofouling layer on the electrode surfaces, as originally discussed in [44, 45]. This mechanism may also account for the absence of correlations between the dimensions and the cellular complexity of the FBR and the deterioration of MEA recording performance over time. The most straightforward conclusion from these findings is that the elimination of microglia adhering to the surfaces of the implanted electrodes should mitigate the problem of MEA performance deterioration.

Using the CSF1R antagonist PLX5622 and Perforated Polyimide-based Multielectrode array Platform (PPMP) that can be sectioned along with the surrounding tissue, we examined this hypothesis in a previous publication [41]. Although ad libitum feeding of adult female rats with PLX5622 eliminated approximately 95% of the cortical microglia and the overall cell composition and dimensions of the encapsulating scar was reduced, the implant recording performance did not improve. This was accounted for by the unexpected finding that whereas 95% of the cortical microglia were eliminated, PLX5622-resistant microglia still adhered to the implant and electrode surfaces [41].

Inspired by the traumatic brain injury literature demonstrating that the recovery of brain parenchyma from mechanical injury is age-dependent [46–48], here for the first time, we examined whether: (a) in young rats, 74±8 gr, 4 weeks old at the onset of the experiments, cortical tissue-recovery following MEA implantation would result in smaller inflammatory scarring as compared to older rats (242 ±18 gr, 9 weeks old at the experimental onset), and (b), in young rats, CSF1R antagonist treatment would eliminate the microglia including the PLX-resistant population that adheres in adults to the MEA surfaces. The findings confirm that the recording performance of MEA was significantly improved in young as compared to adult rats and that microglia elimination further improved recording performance. The temporal correlations between the PPMP implant performance and immunohistological observations as a function of time thus support the hypothesis that microglia adhering to the implant surfaces constitute a major underlying cause of PPMP performance deterioration over time. In addition, the results suggest that young rats are an advantageous model to study months-long, electrophysiological/behavioral mechanisms.

## 2. Method

### 2.1 Animals

All the used procedures were approved by the Committee for Animal Experimentation at the Institute of Life Sciences of the Hebrew University of Jerusalem (accredited by the Association for Assessment and Accreditation of Laboratory Animal Care). The immunohistological and functional recordings were made on young female Sprague Dawley rats, 74±8 gr, 4 weeks old at the onset of the experiments, and adult female Sprague Dawley rats, 242 ±18 gr, 9 weeks old at the onset of the experiments. For microglial ablation, the rats were fed ad libitum with the CSF1R antagonists PLX5622 or PLX3397. 1200 mg/Kg-food PLX5622, (Plexxikon Inc., Berkeley, USA), and 1000 mg/Kg-food PLX3397 (Hy-16749, Med ChemExpress). The control diet consisted of the same base formula. PLX5622 was provided by Plexxikon Inc. and was formulated in AIN-76A standard chow by Research Diets Inc. The PLX3397 was dissolved in dimethyl sulfoxide (DMSO ;Sigma-Aldrich, catalog #276855). The solution was added to Teklad Global 18% Protein Rodent Diet (2918, Envigo) as previously described [49].

### 2.2 Neuroimplants

For the current study we used in-house fabricated Perforated Polyimide (PI) based MEA Platforms (PPMPs, Fig. 1) as detailed earlier by our laboratory [20]. Two types of single shaft, 280 μm wide, 16 μm thick platforms carrying 15 planar gold electrodes were used: (1) free floating (untethered) and fixed-to-the-skull (tethered) nonfunctional (dummy) platforms for the immunohistological studies, and (2), functional platforms for electrophysiological recordings [34]. These platforms were implanted into the motor cortex as detailed in our earlier publications [20] with slightly modified platform implantation coordinates: for the immunohistology examinations, the dummy electrodes were implanted as follows: in adult rats: AP= 3.5, ML= +2.5 mm, and DV= -1.1.8 mm and in young rats: AP= 3.4 mm, ML= 2.5 mm, and DV= -1.8 mm. To optimize the FP recordings, the functional electrodes were implanted in adult rats at AP= 4.2-4.3 mm, ML= 3.4-4.1 mm, (selected during the surgery according to morphology of the skull) and DV= -1.27--1.57 mm and in young rats at: AP= 4 mm, ML= 2.6-3.7 mm, and DV= -1.41--1.82 mm.

**Fig. 1.**
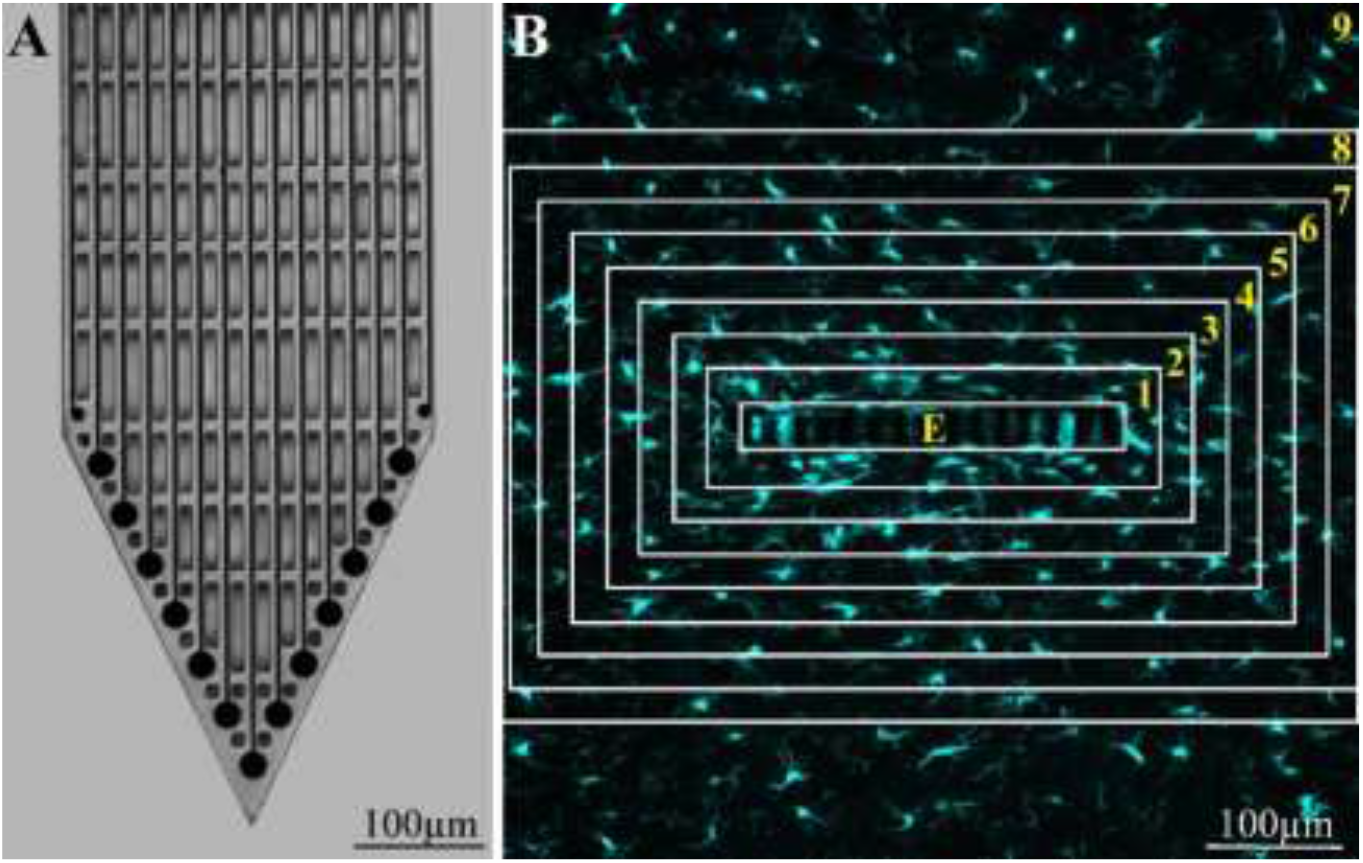
(A) For orientation purposes, a light microscope micrograph of the perforated polyimide based MEA platform (PPMP) tip and a confocal microscope image of a horizontal slice through the PPMP (E) along with the cortical parenchyma around it (B). (A) Shown are 13 large and 2 small planar electrodes (dark circles) and the perforated microarchitecture of the PPMP’s distal part. (B) The cell densities or fluorescent intensity images of different cell types within the implant pores (E-central rectangle; see enlargements in Figures 4 and 5) and within 25 μm wide centripetal shells around it were counted or measured at different points in time post-PPMP implantation and processed to establish the density or normalized fluorescent intensity level (NFI) at a given distance around the electrode. The image in (B) depicts an example of microglia labelled by Iba-1.

### 2.3 Tissue processing for immunohistology

Rats implanted with tethered or “floating” dummy MEA platforms were sacrificed at 3 days, 1, 2, or 4 weeks after implantation. Brain fixation, cryo-sectioning and immunohistological labeling of microglia, astrocytes, and neurons were conducted as detailed in our previous publications [20, 41, 42].

### 2.4 Microscopy

Confocal image stacks of the immunolabeled slices were acquired with an Olympus FLUOVIEW FV3000 confocal scan head coupled to an IX83 inverted microscope, using a 20X air objective (NA = 0.75) and 60X oil objective (NA=1.4). Scanning was conducted as previously described [20, 41, 42]. Typically, 15-30 confocal slices were acquired from the perforated segment of the implant, with a vertical spacing of 1μm (Fig. 1).

### 2.5 Electrophysiology

Electrophysiological experiments were conducted on the control-treated adult rats, control, and CSF1R antagonist-treated young rats. In these experiments, the microglia were eliminated by PLX3397 (Pexidartinib). PLX3397, like PLX5622, eliminates over 90-95% of the microglia within 5-7 days of feeding [41, 50, 51]. Since PLX3397 was approved by the FDA for treatments of adult patients with symptomatic tenosynovial giant cell tumor (TGCT), its intrinsic advantage over PLX5622 lies in its potential medical applications [52, 53].

Electrophysiological recordings were obtained weekly, starting on the day of surgery up to 6 weeks post-implantation. In total 15 rats were recorded from, 5 adults, 6 young controls, and 4 PLX3397 treated young rats. During each recording session, the rats were placed in an induction chamber and anesthetized by exposure to 8% sevoflurane (Piramal Critical Care Inc., Bethlehem, PA, United States) using the SomnoSuite Low-Flow Anesthesia System (Kent Scientific Corporation, Torrington, CT, United States). When the breathing rate slowed down, the rats were placed in a stereotactic frame with exposure to 4% sevoflurane and were connected to a nanoZ impedance tester (Plexon). The total in-vivo impedance was measured at 1 KHz, by passing 1.4 nA through each of the functional electrodes across the surrounding parenchyma and the bone screw to the ground. Thereafter the connections were switched to a 16-channel multichannel system wireless amplifier (W2100-HS16, Multi Channels systems, a division of Harvard Bioscience, Inc.) and electrophysiological recordings were collected in awake rats moving freely in a large container for 10 minutes. The signals were then amplified and digitized. As a ground reference, we used the screw attaching the cement structure and the MEA platform to the skull (Precision Technology Supplies Ltd., United Kingdom). At the onset of each experiment, the impedances were measured before surgery and electrodes with distinctively high impedances of > 6 MΩ at 1 kHz were excluded from the analysis. During each recording session, electrophysiological data were collected at a 20 kHz sampling rate, and a bandpass 300-3000 Hz filter was applied for single-unit recordings. Spike sorting was performed using the fully automatic spike-sorting implementation described by Chaure et al. [54]. The minimum threshold for spike detection was set to both negative and positive 3.5 times the standard deviation (SD) of the background channel noise. The background channel noise was defined as twice the median (raw voltage)/0.6745 [55, 56]. The spike time window was set to 20 samples (1 ms) before and 44 samples (2.2 ms) after each threshold crossing event and the spike traces were automatically sorted into clusters. The minimal cluster size was set to 20 spikes. Clusters with obvious artifact wave features such as very high amplitudes (>30 times background noise amplitude) and similar negative and positive amplitude phases, were removed. These artifacts were assumed to be due to the sudden appearance of very high voltage amplitudes simultaneously in most/all channels caused by rat head movements or the noise from surrounding devices during the recording session. In cases where identical spike shapes were wrongly separated into different clusters because of brain micro-movements that affected the amplitude or because of spike jittering, the clusters were either manually merged or the cluster with the higher number of spikes was selected. Signal amplitudes were calculated as the peak-to-peak amplitude of the mean waveform of each cluster. The signal-to-noise ratio (SNR) for each cluster was calculated as the average spike amplitude divided by the background noise of each channel. The percentage of channels detecting spikes in a given recording session was calculated as the ratio of the total number of channels detecting spikes to the total number of functional electrodes.

### 2.6 Image processing, analysis and statistics

The image processing was implemented using the Fiji distribution of ImageJ [57, 58], as described previously [20]. Two methods of analysis and representation of the cell densities in contact and around the PPMPs were used: (1) The densities of the astrocytes and neurons, including their cell bodies and neurites, were analyzed and depicted as their relative fluorescent intensities with respect to the normal background; henceforth, the Normalized Fluorescent Intensities (NFI) values. (2) The NFI approach could not be applied to compare the microglia densities between the control rats and the PLX5622/PLX3397 treated rats since the PLX5622/PLX3397 treatment reduced the cortical microglia densities to approximately 5% as compared to the controls. Specifically, the denominators to calculate the NFI (fluorescent intensity near the implant/ fluorescent intensity remote from the implant) of the control and PLX5622/PLX3397 treated rats could not be compared. For this reason, the density values of the microglia (for both the control experiments and the PLX5622/PLX3397 treated rats) were calculated using automated counting, which was manually confirmed for accuracy. The microglia densities are thus expressed as the average number of cells per mm^2^ in a given shell. To count the microglia, we used merged images of Iba1 labelled cells complemented by DAPI labeling. The microglia nuclei were unequivocally identified using the characteristic heterochromatin distribution, nuclear shape, and size [20]. The neuron densities in time and space are given in both NFI units and number /mm^2^. Because of the relatively delicate cytoplasm of the astrocytes and their elaborate branching pattern, the automatic counting of astrocytes (GFAP co-labeled with DAPI) resulted in unacceptable mismatching with the manual counts. Therefore, the astrocyte densities are presented here in NFI values alone.

The average fluorescent values and cell counts characterizing the FBR in space and time were measured and calculated from cortical brain slices prepared from 2-10 hemispheres/experimental points in time. Each brain hemisphere was used to prepare 1 to 6 tissue slices from the porous segment of the platform (Fig. 1). Each slice was used to prepare a single maximal projected image generated by 10 consecutive optical sections. The sample sizes of the immunohistological sections of the nonfunctional PPMPs are listed in Supplementary Table S1. Differences between the cell density values at different distances from the implant and at different points in time were assessed by a t-test for two samples assuming unequal variances. For all tests, a p value < 0.01 indicates a statistically significant difference (Supplementary Table S2).

## 3. Results

To characterize the neuro-inflammatory processes induced by PPMP implantation in young and adult female rat cortices, and compare the effects of microglia elimination by ad libitum CSF1R antagonist feeding (PLX5622 or PLX3397) we: (1) mapped the microglia, astrocyte and neuron densities in intact cortices (not implanted with MEA platforms) of young and adult rats, (2) mapped and compared the spatiotemporal distribution patterns of these cell types in control young and adult rats following the implantation of untethered and tethered-PPMP and (3), examined the effects of ad libitum PLX5622 diets from the 4^th^ day post-PPMP implantation on inflammatory processes in young and adult rats. Note that our previously reported observations [41] showed that the spatiotemporal distribution of microglia in adult female rats fed with PLX5622 chow for a week before PPMP implantation and onward, or 4 days after implantations and onward, did not differ when imaged one to eight weeks post-PPMP implantation. For this analysis, we examined immunohistological sections of the cortical parenchyma along with the implants at 3 days, 1, 2 and 4 weeks post-PPMP implantation. These points in time were selected based on our earlier studies indicating that in adult female rats, these junctures represent discernible stages of the implant-induced inflammatory cascades [20, 41].

Recall that the inhibition of the CSF1R in adult mice and rats by either PLX3397 or PLX5622 chow leads to the elimination of 90-95% of the microglia within 5-7 days of feeding [41, 50, 51] and that approximately 5% of the cortical microglia population in mice and rats resist CSF1R inhibition [41, 59, 60]. As a reference for the quantitative assessments of cell densities, we established that the average microglia density in intact young and adult female rats was 363±39 cells/mm^2^ and 263±34 cells/mm^2^, respectively. The average cortical neuron densities were 1611±223 cells/mm^2^ in the young and 1106±179 cells/mm^2^ in the adult rats, respectively. Because of the small size of the astrocyte cell bodies cytoplasm and the elaborate branching patterns, automated quantifying the cell bodies densities (number/mm^2^) was imprecise (see methods). For this reason, the astrocyte spatiotemporal distribution is expressed in what follows in NFI values. The differences in the spatiotemporal dynamics of microglia, astrocytes, and neurons in the control and the PLX5622 treated young and adult rats are described below.

### 3.1 Microglia: inflammatory cascades after PPMP implantation in the control and CSF1R treated young and adult rats

In an earlier study conducted in our laboratory, we reported that PPMP implantation into control adult female rat cortices was followed by the migration of microglia to occupy the PPMP pores, adhesion of the microglia to the implant and electrode surfaces, proliferation of the microglia in the vicinity of the implant and increases in microglia density in shells 1-5 around the implants (0-125 μm, Figs. 2, 3, S1 and S2). The increased microglia density spontaneously subsided with time in these shells but microglia remained attached to the platform and the first shell around it (0-25μm) for over 8 weeks post-PPMP implantation [20]. Buttressed by electrophysiological examination, we then proposed that the microglia adhering to the PPMPs surface electrically insulate the electrodes from the surrounding tissue.

**Fig. 2.**
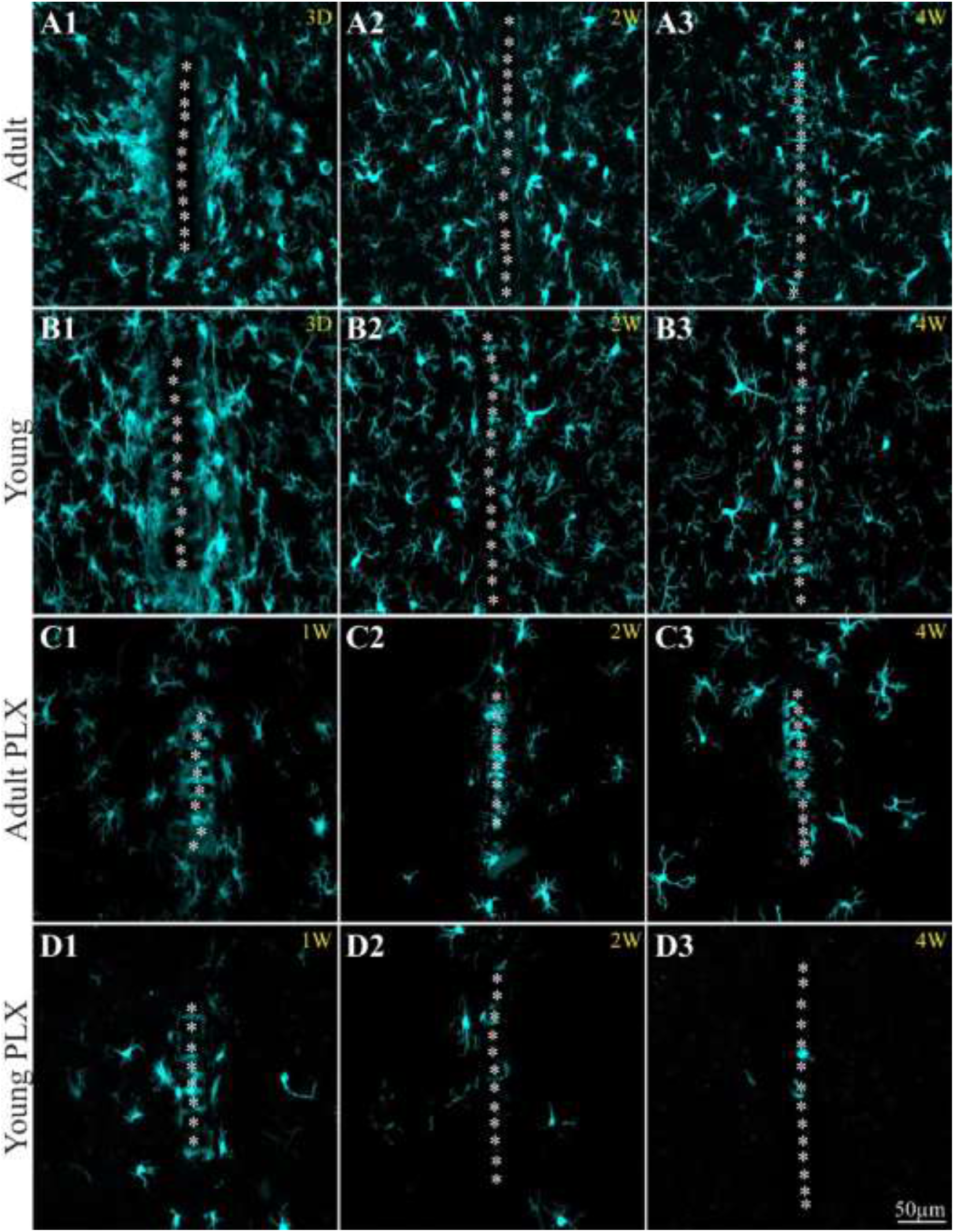
Comparison of the spatiotemporal distribution of the microglia in contact and around implanted PPMPs in the control and PLX5622 treated adult and young rats. The microglia were labelled by the Iba-1 antibody. For orientation, the white asterisks show the solid polyimide “ridges” between the pores of the PPMP. (A1-A3) control adults, on day 3, weeks 2 and 4 after PPMP implantation. (B1-B3) control young rats on day 3, weeks 2 and 4 post PPMP implantation. (C1-C3) PLX5622 treated adults, (D1-D3) PLX5622 treated young rats, 1, 2 and 4 weeks post-PPMP implantation. Note that the microglia densities within, in contact, and around the implant are larger in the adult (A1-A3) than in the young rats (B1-B3, for quantitative data see Figure 3). Ad libitum feeding of the adult and young rats with PLX5622 (starting on the fourth day post PPMP implantation) greatly reduced the microglia densities around the implants. Unlike in young rats where the density of the PLX5622-resistant microglia was minimal (D2-D3, and see also supplementary Figure 1), in the adults, denser PLX5622-resistant microglia remained attached to the platform (C2-C3).

**Fig. 3.**
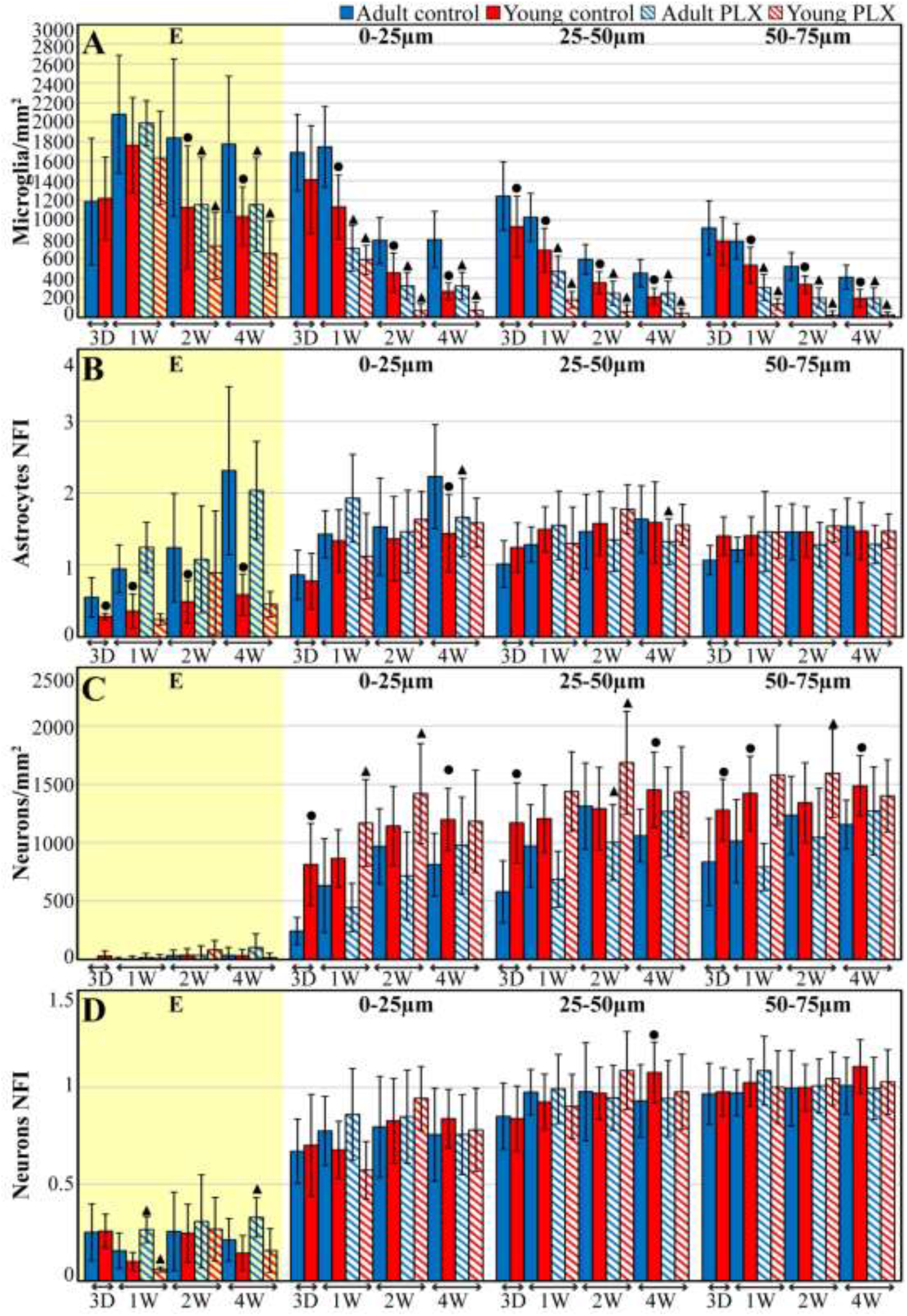
The average densities of the microglia (A), normalized fluorescent intensity (NFI) of the astrocytes (B), neuron density and NFI (C and D respectively) within the PPMP (yellow background, E), in contact (shell 1-0-25μm) and around implanted PPMPs (shells 2 and 3, 25-50 and 50-75μm) in the control and PLX5622 treated adult and young rats at different points in time post-PPMP implantation. Solid blue columns depict control adult cortices. Solid red columns depict control young rats. Diagonal blue and red stripes depict cortices of adult and young rats fed with PLX5622 4 days after PPMP implantation and thereafter. The times post- platform implantation (3 days, 1, 2, and 4 weeks) and the distance of the measured averages are given below the X axis of the histograms. Vertical lines correspond to one standard deviation. Black circles indicate significant differences of P<0.01 between the value of the control adults and the control young rats; triangles indicate significant differences of P<0.01 between PLX5622 treated adults or young rats and the control adults or young rats. Similar data depicting the spatiotemporal densities and NFI values over a centripetal distance of 300μm are shown in Fig. S2.

In the young implanted control rats, the average microglia density was smaller than that recorded in adults from day 3 post implantation and onwards, within the implant pores, attached to the implant surfaces and electrodes, as well as in shells 1-6 around it (Figs. 2, 3, S1 and S2). The significant difference (P ≤ 0.01) in microglia densities between implanted young and adult rats persisted over the 4 weeks of observations and over a centripetal distance of 0-100 μm from the PPMP surfaces (1-4^th^ shells, Figs. 2, 3 and S2). If the hypothesis assuming that microglia adhering to the PPMP surfaces, rather than the multicellular encapsulating inflammatory scar, is correct, than this observation would imply that the recording performance of PPMPs implanted into young rat’s cortices may be better than that of adults.

### 3.2 PLX5622 treated implanted young versus adult rats

The spatiotemporal distribution dynamics of microglia after PPMP implantation in cortices of the young rats fed with PLX5622 chow from the fourth day post-PPMP implantation and onward differed in a number of ways from the PLX5622 -treated adult rats described previously by our laboratory [41] and here. In both the PPMP-implanted adults and young rats, PLX5622 feeding led within a week to the elimination of approximately 85% of the cortical microglia, which rose to ~95% on day 12 post-implantation. In both the young and adult rats fed by PLX5622 diet, a PLX-resistant subpopulation was observed within the first shell around the implant and attached to it (Figs. 2, 3, S1, S2 and [41]). Whereas in the adult rats the PLX5622-resistant microglia adhered to the implant surface and thus appeared to insulate the MEA from its surrounding tissue (as detailed in a previous study conducted in our laboratory [41] and shown in Figs. 2C and S1), in young rats, only a small number of microglia remained adhering to parts of the implant surfaces (Figs. 2D and S1). This observation may imply that the recording performance of PPMPs implanted into young rats fed by PLX5622 should be somewhat better than that of control young rats, and significantly better than in adults.

To better resolve the structural interfaces formed between the microglia and the implant we used a high magnification (60X), high resolution (numerical aperture 1.4) objective and merged light microscope images of the implants with confocal images of the microglia (Figs. 4 and 5). Using this setting revealed that even in implanted control young rats in which the PPMP appeared to be almost “microglia free” under standard resolution confocal microscopy (Figs. 2, 3 and S1), delicate microglia branches were seen to adhere to the PPMP surfaces (Fig. 4) and interpose between the platform and the neurons. These confocal images were consistent with our earlier transmission electron microscope observations showing that delicate microglia branches adhere tightly to the PPMP surfaces [42]. Theoretically, these adhering branches suffice to electrically insulate the electrodes from the surroundings neurons [20, 41, 42]. Importantly, in young rats fed with PLX5622 in which the presence of microglia in contact with the implant was further reduced, microglia-free surfaces were often imaged (Fig. 5). Thus, as suggested previously and further discussed here, the tight adhesion of microglia branches to the PPMPs may play a critical role in insulating the implanted electrodes from the surrounding neurons [20, 41–43].

**Fig. 4.**
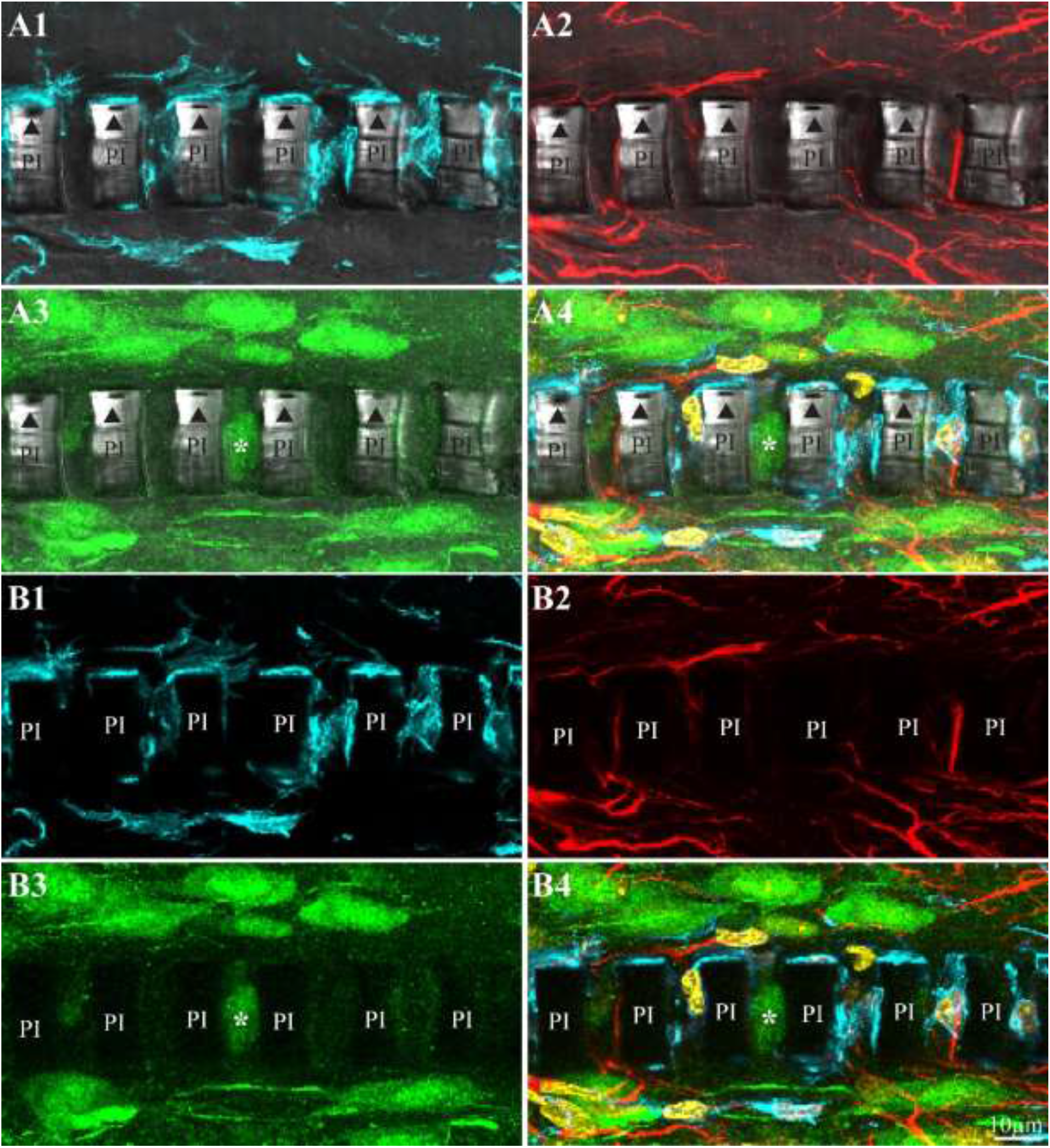
High resolution, large magnification (X60 objective, numerical aperture 1.4) confocal microscope images illustrating the interfaces formed between the implanted PPMPs and the cells around it. Control young rat, 2 weeks after implantation. Images A1-A4 were prepared by merging light microscope images of the implanted polyimide platforms and the corresponding confocal images of the parenchyma surrounding them. (B1-B4) Confocal images of the parenchyma as in (A) without the polyimide platform light microscope image (A1&B1) Iba-1 antibody labeled microglia (cyan), (A2&B2) GFAP antibody labeled astrocytes (red), (A3&B3) NeuN and NF antibodies labeled neurons (green), (A4&B4) merged image of microglia, astrocytes, neurons and DAPI labeled nuclei (yellow). Note that the thin microglia cytoplasm adhering to the polyimide surface extends into the pores of the platform and interposes between the platform and neurons. In (A3) a neuronal cell body is seen to reside between two PI ridges. PI-polyimide “ridges”, triangles-conducting lines, asterisks indicate a neuronal cell body residing within a platform pore.

**Fig. 5.**
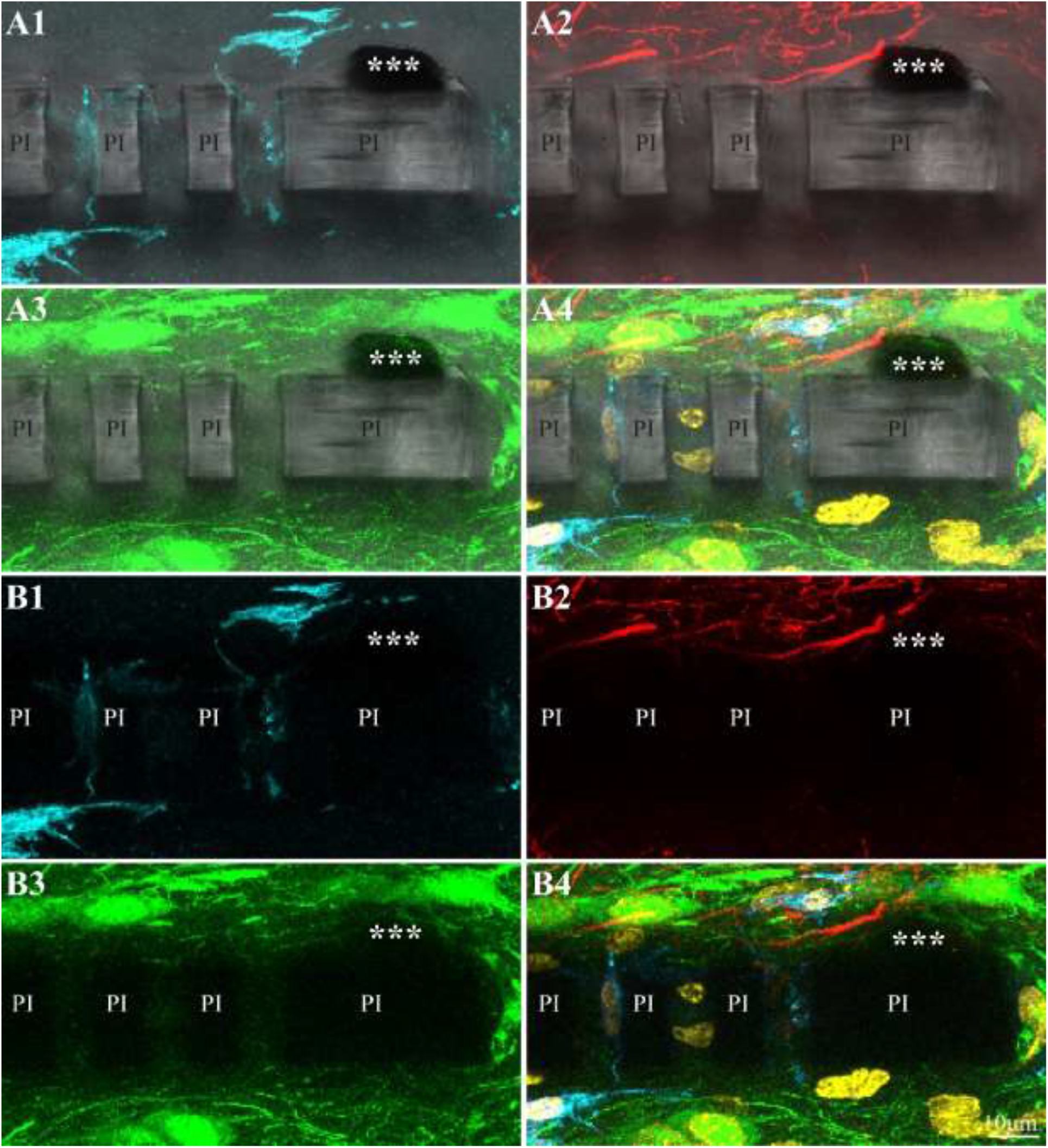
High resolution, large magnification (X60 objective, numerical aperture 1.4) confocal microscope images illustrating the interfaces formed between implanted PPMPs and the cells around it. In a PLX5622 treated young rat (4 weeks after implantation). Images A1-A4 were prepared by merging light microscope images of the implanted polyimide platforms and the corresponding confocal images of the parenchyma surrounding them. (B1-B4) Confocal images of the parenchyma as in (A) without the light microscope image. (A1&B1) Iba-1 antibody labeled microglia (cyan), (A2&B2) GFAP antibody labeled astrocytes (red), (A3&B3) NeuN and NF antibody labeled neurons (green), (A4&B4) merged image of microglia, astrocytes, neurons and DAPI labeled nuclei (yellow). Note that in the PLX5622 treated young rats, the surface of the platform is almost entirely free of microglia. PI - polyimide “ridges”; three asterisks indicate a section through an electrode.

By co-labelling the Iba-1 positive PLX5622-insensitive microglia of the young implanted rats with anti-rat TMEM119 that identifies microglia-specific transmembrane proteins, we confirmed that (as in adults [41]) the cells adhering to the implants were genuine microglia and not other Iba-1 positive cell types that could have infiltrated the brain tissue through the damaged surroundings of the implanted platform (Fig. S3A). Furthermore, as was the case in the adult rats, co-labelling of the PLX5622-insensitive microglia by Iba-1 and KI67 (which labels cells undergoing mitosis) established that a fraction of the microglia observed in the implant vicinity underwent proliferation (Fig. S3B).

It is well-established that cortical implants fixed to the skull (tethered implants), a setting that better imitates the configuration of functional platforms, induces a more intensive FBR than free floating implants. In a previous study we found that tethering dummy PPMPs to the skull of adult rats did not worsen the FBR in general or the microglia densities in particular as compared to untethered dummy PPMPs [20]. Here we observed that in the control and PLX5622 fed young rats, tethered PPMP led to increased microglia densities within the implant pores as in the adult rats but to a smaller extent in the first shell around the implant and farther away from it (Figs. S4 and S5).

In summary, after an initial surge of increased microglia densities after PPMP implantation, the microglia densities in both adult and young rats underwent gradual recovery in the shells around the implant from the second week of implantation onward (Figs. 2 and 3). The recovery rate (reduced density) was faster in young rats than in adults. PLX5622 treatment facilitated the rate and extent of reduction of microglia densities around the implants. Importantly, whereas in the control and the PLX5622-implanted adult rats the (PLX resistant) microglia continued to adhere to the implant surface for over 8 weeks [41], in the control implanted young rats the microglia density in contact with the implant declined significantly as compared to the adult rats. Subsequent to PLX5622 treatment the microglia were almost entirely eliminated from untethered implant surfaces.

### 3.3 Astrocytes: activation in control and PLX5622 treated young and adult rats

Note that NFI values are used in this section to compare the relative change in astrocyte density within (but not between) the adult or young age groups. In both the control implanted young and adult rats, activation of astrocytes in the form of branch outgrowths and cell proliferation after PPMP implantation was delayed as compared to the microglia by approximately one week [20, 41]. Thereafter, whereas in adults, the astrocyte NFI values increased within the implant and the first shell around it by 24-53% (to 1.24±0.75 within the implant and to 1.53±0.68 in the first shell) on the second week post-implantation, and up to 123-131% (2.31±1.17 within the implant and 2.23±0.72 in the first shell) on the fourth week postimplantation (Fig. 3 and Supplementary Table 2), in young rats the increase in NFI value was significantly less than one (P=4.8*10^-^ for 2 weeks and 2.8*10^-^ for 4 weeks) within the implant pores and 36-44% (1.36±0.59 on the second and 1.44±0.54 on the fourth week postimplantation) in the first shell around it (Figs. 3B, 4A2 & B2 and Supplementary Table 2). The astrocyte NFI values of both the young and adult rats diminished with distance from the implant surface but remained >1 even at a distance of 200μm from the PPMPs for over a period of > 4 weeks post-implantation (Fig. 3B and S2B). PLX5622 treatment had no significant effect on the astrocyte NFI values after PPMP implantation into adult and young cortices. Importantly, the astrocyte NFI values in the first shell increased in young rats when tethering the electrodes to the skull by a factor of 1.39-1.57 as compared to the untethered configuration in PLX5622 fed and in the control rats, respectively (Figs. S4B, S5 and Supplementary Table 2). High magnification confocal images of the astrocytes in the control and PLX5622 treated young rats revealed that delicate astrocyte branches loosely ramified near the PPMP surface (Figs. 4 and 5).

Taken together, whereas in PPMP implantation into young and adult rats, the microglia densities decreased continuously from the first/second week and onward (Figs. 2 and 3), the astrocyte NFI levels continued to increase from the first week post-implantation and onward, reaching NFI values of >2 in the adults and >1.4 in the young rats in the fourth week (see also NFIs in [20, 41]) and in the first shell around it (Fig. 3, 0-25μm and 6). The loose astrocyte branches observed in between the neurons and the implanted MEA might have contributed slightly to the impedance between the electrodes and the neurons [20].

### 3.4 Neurons: density recovery in the control and PLX5622 treated young and adult rats

Given the significant differences in the microglia and astrocyte spatiotemporal distribution patterns between young and adult rats and the age-related differences in the responses to PLX5622 treatment, we next examined whether these alterations were associated with differences in the spatiotemporal distribution of the neurons around and in contact with the implant.

This was analyzed in two ways: (1) the density of neuronal cell bodies was quantitatively mapped using the NeuN labelling (Fig. 3C); (2) The spatiotemporal distribution of the neurons including axons, dendrites and cell bodies as labelled by NeuN and NF were mapped using the NFI protocols (Fig. 3D, S2D and Supplementary Table 2 and see [20, 41]).

In both young and adult rats, neuronal cell bodies were occasionally seen to occupy the PPMP pores (for example Fig. 4B3). Typically, low average neuronal cell body counts of <38 cell bodies/mm^2^, and low average NFI values of <0.33 were observed within the PPMP over the 4-week period of observations. The low fluorescent level detected within the PPMPs reflects the outgrowth of neurites along the platform’s surface and into the platform’s perforations, and the auto-fluorescence of the PI (Fig. 3D and S2D).

The damage caused by PPMP implantation to the neuron densities along the insertion path was similar in young and adult rats. Decreased neuron counts were recorded on the third day post-insertion in the first, second and third shells around the implant (0-75μm, Figs. 3C S2C). In adults, young controls, and the PLX5622-treated individuals, the average neuron NFI values (including the neurites and neuron somata), gradually increased, reaching 76-84% of the control levels in the 4^th^ week post-implantation in shell one (0-25μm, Figs. 3 and 6 C, D). In adults, the cell body density recovered to 814±268 cells/mm in the first shell and to 1063±228 and 1168±233 cells/mm^2^ in the second and third shells in the 4^th^ week post-implantation. In the control young rats, recovery reached higher values of 1243±313 cells/mm^2^ in the first shell and 1464±341/ 1490±258 cells/mm^2^ in shells 2-3 (25-75μm). These differences do not reflect a faster recovery rate but rather are related to the differences in the control density of the neurons in young and adult rats (1611±223 cells/mm^2^ in young and 1106±179 cells/mm^2^ in adult rats). The PLX5622 treatment of the young rats improved the initial recovery rate of neuron densities (as compared to the control young rats) to 1169±372 cells/mm^2^ in the first shell in the first week post-implantation and to 1454±434cells/mm^2^, 1718±423 and 1640±328 cells/mm^2^ in the first, second and third shells in the second week post-implantation (Fig. 3C).

**Fig. 6.**
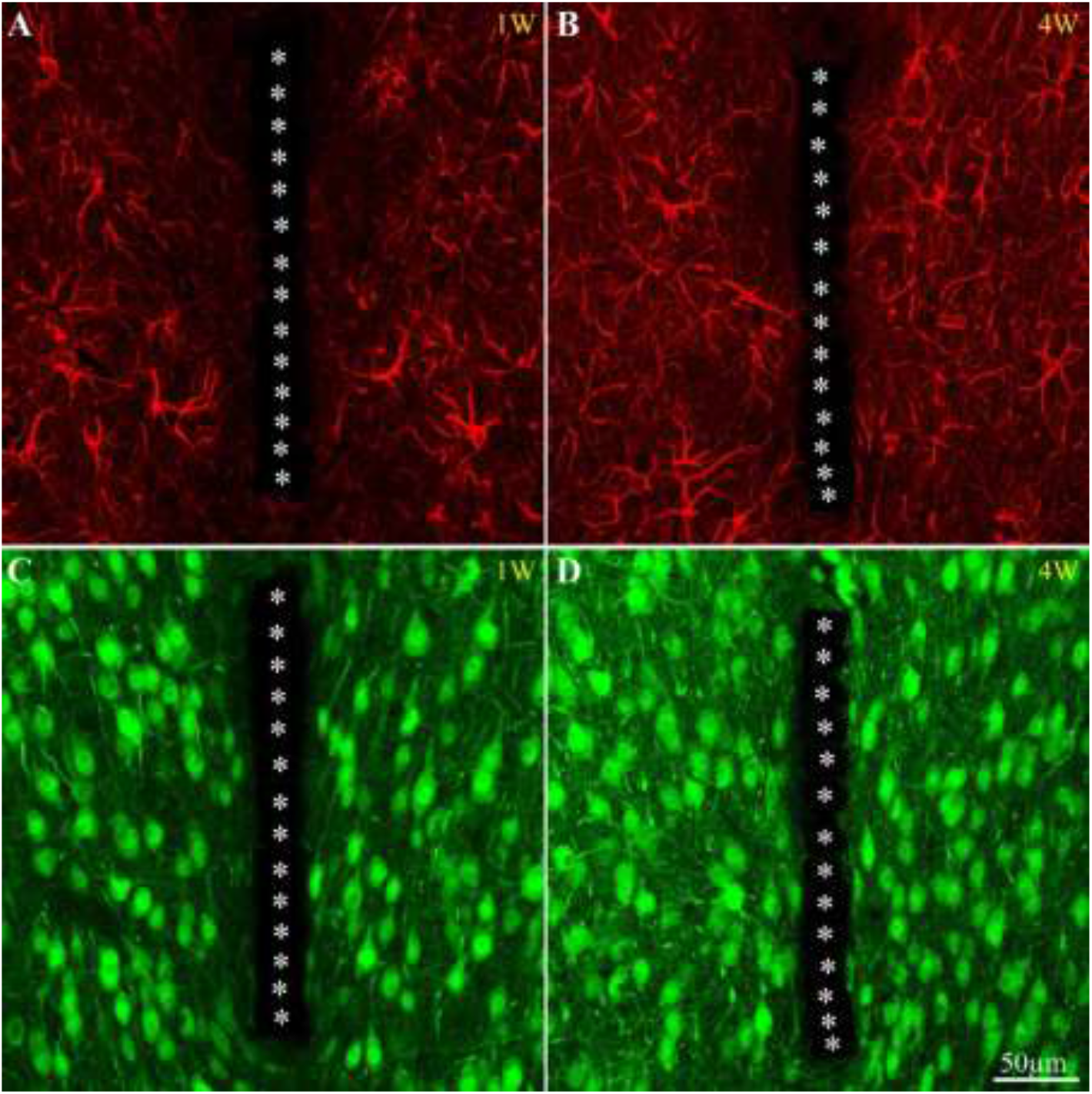
Examples of astrocyte and neuron distributions around the implanted PPMP of young rat cortices fed with PLX5622 chow from the fourth day post implantation onward. (A, B) Astrocytes, (C, D), neurons. (A) and (C), a week, (B) and (D) four weeks, after PPMP implantation. For orientation, the white asterisks indicate the solid PI “ridges” between the pores of the PPMP. Note the density of neuronal cell bodies within a distance of less than 150μm from the PPMP platform surface.

Comparison of the neuronal somata densities and the NFI values of the tethered and free-floating PPMPs did not reveal significant differences (Fig. S4).

Thus overall, the neuronal regenerative rates following PPMP implantation were similar in young and adult rats. Treatment of young rats with PLX5622 further enhanced the initial recovery and reduced the immunohistological footprint of the inflammatory processes caused by the implantation (Figs. 3C and 6 C&D). Consistent with our previous studies, it was found that neuronal cell bodies reside at a distance of <10 μm from the implanted MEA surfaces (Figs. 4, 5, 6 and [42]).

### 3.5 PPMP recording performance, control vs. PLX3397 treated adult and young rats

Next, we examined whether the differences in the inflammatory encapsulation characteristics described above in the control implanted adult, control young rats, and CSF1R antagonist-treated young rats would be expressed in terms of recording performance. We thus assessed the percentage of operating platforms, the percentage of viable electrodes/PPMP, FP amplitudes and the signal to noise level. To that end, we recorded spontaneous FPs from freely behaving rats during PPMP implantation and once a week for 6 weeks (for example Fig. 7). In this series of electrophysiological experiments, we eliminated the microglia of young implanted rats using the CSF1R antagonist PLX3397 (Pexidartinib). Since PLX5622 and PLX3397 eliminate over 90-95% of the microglia within 5-7 days of feeding [41, 50, 51], and because PLX3397 was approved by the FDA for the treatment of adult patients with symptomatic tenosynovial giant cell tumor (TGCT) it had the further advantage over PLX5622 of having potential medical applications [52, 53].

**Fig. 7.**
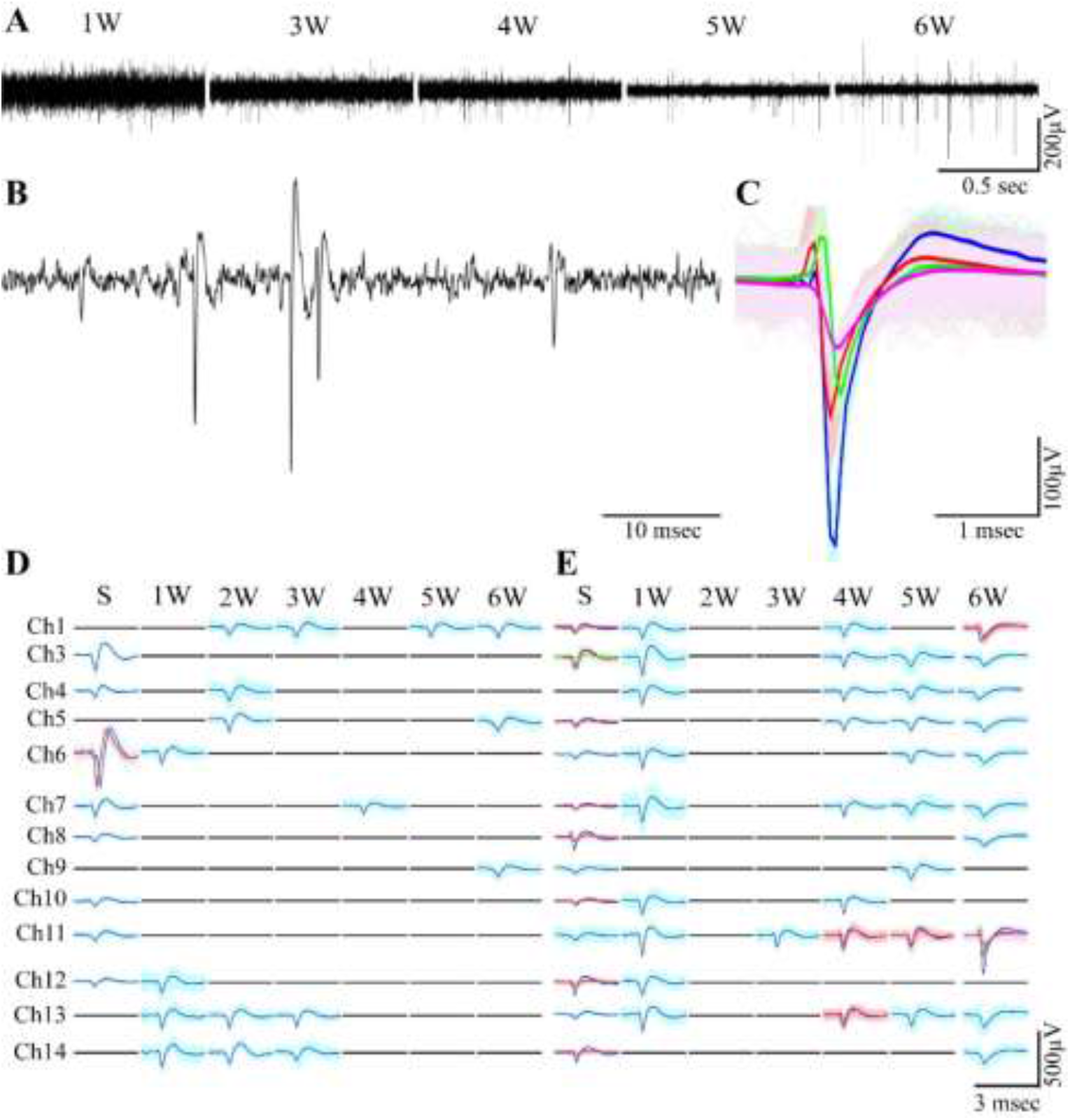
Examples of the time course of spontaneous FP traces and clustered spike shapes recorded by PPMPs for 6 weeks after implantation. (A) An example of voltage traces recorded by one microelectrode from a single PLX-treated young rat for 6 consecutive weeks. (B) Enlarged voltage traces recorded on the 6^th^ week after implantation. The recorded field potentials were sorted by a fully automatic algorithm into (C) spike clusters shown by different colors where the bold line is the average and the spike shape traces are the bright background. An example of the FP clusters recorded throughout the entire 6-week period (D) control and (E) PLX3397 fed young rats.

The FPs amplitudes recorded during the PPMP insertion sessions were similar in adult and young rats (150.8± 63.5 μV in adults and 163.3 ±111.3 μV in young rats, averaging 157.4± 88.7 μV). The average spontaneous FP amplitudes recorded from the control adults, control and PLX3397 fed young rats thereafter were 153.8±37 μV in adults, 151.5±41.3 μV in young rats and 166.1±54 in young rats fed by PLX3397 (for an example see Fig. 7; in total 15 rats, 5 adults, 6 young controls, and 4 PLX3397 treated). The average noise level of the implanted PPMPs on the day of implantation was 22±4.8 μV. Thereafter, the noise levels of the live electrodes increased in all three experimental groups reaching maxima on the first or second week after implantation and then gradually declined (Fig. 7). The maximal noise levels were 36.3±5.5 μV, 32±6 μV and 35.8±7.2 μV in the control adult, control young and PLX treated rats respectively. Aligned with the differences in noise levels, the signal to noise ratio ranged from 4 to 17.5.

The differences in recording performance among the experimental groups are depicted in Fig. 8 (and Fig. S5) in which the percentage of active PPMPs and live electrodes over a six-week period are compared. An operating platform was defined here as a platform in which at least one electrode was recording FPs (live electrode). The percentage of recording electrodes in a given session was calculated as the ratio of the number of electrodes that actually recorded spontaneous FPs to the number of electrodes that were capable of recording (that were not damaged during the fabrication process).

**Fig. 8.**
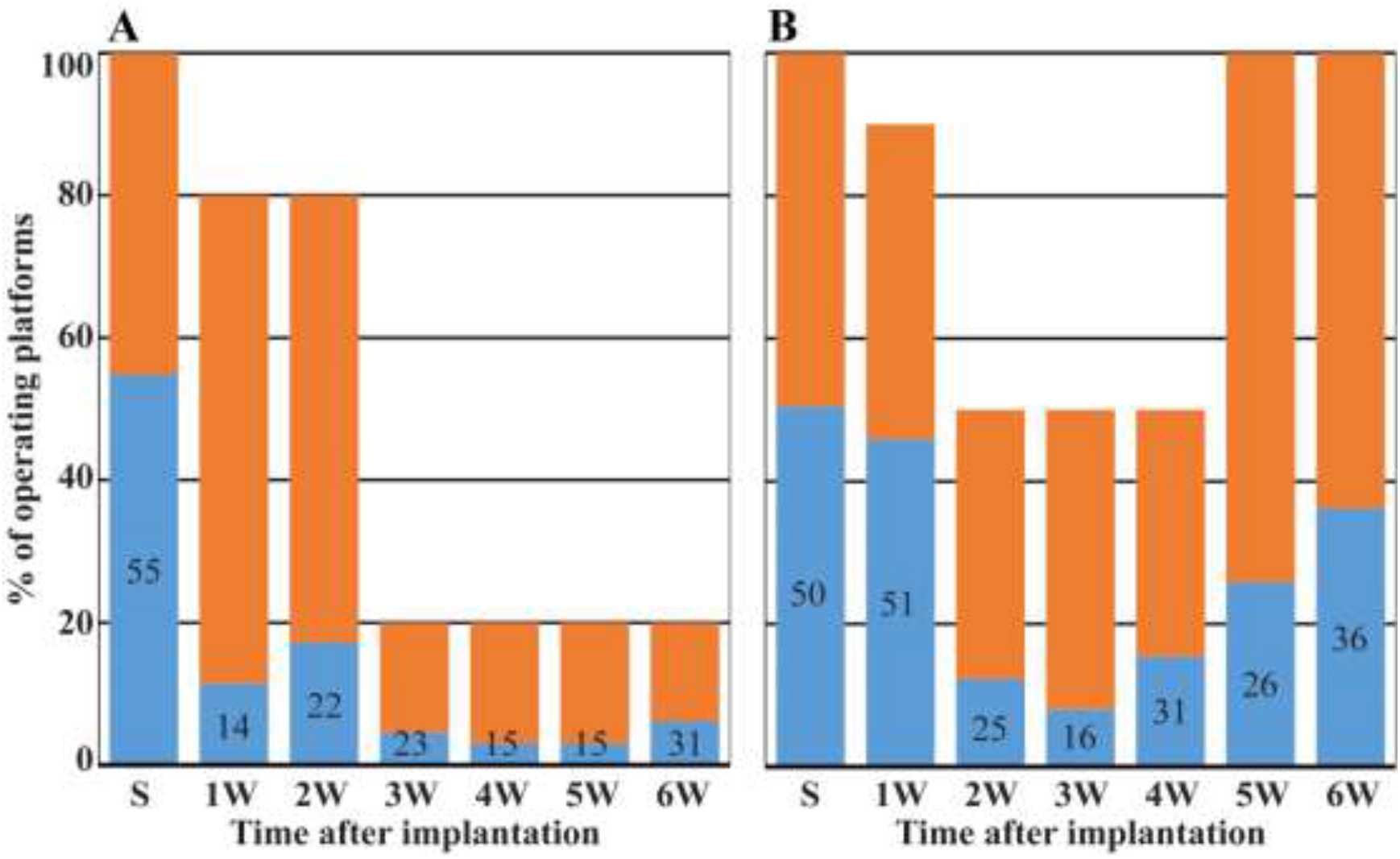
Operating PPMPs and live electrodes in control adults (A), control and PLX3397 fed young rats (B) as a function of time post PPMP implantation. In orange, percentage of operating platforms, in blue, the percentage of live electrodes. In adult rats, the percentage of operating platforms and live electrodes declined with time and did not recover (A). In contrast, in control young rats and rats treated with PLX3397 (B) the percentage of operating platforms and live electrodes began to spontaneously recover towards the values recorded on the day of implantation. Separate histograms for young control and PLX-treated young rats are presented in Supplementary Fig. S4.

In the control adults, the percentage of functional PPMPs deteriorated from 100% on the day of implantation to 20% on the sixth week after implantation (Fig. 8A). This deterioration was associated with a decline in the percentage of live electrodes from 55% to 31%. These deterioration trends are consistent with our earlier studies conducted on adult control rats, and adult rats treated with PLX5622 [20, 41]. Recall that in both groups, microglia continued to adhere to the implanted surfaces. In contrast, in young controls and PLX5622 treated young rats, the surfaces of the implants were almost free of adhering microglia (Figs. 2, 3, 4 and 5), but in both control and PLX fed young rats tethered PPMP elevated the microglia densities within the implant pores (Figs. S4 and S5).

To emphasize these different performance trends in relation to the microglia distribution pattern, Fig. 8 presents the deterioration trends of the young control and PLX3397 treated young rats together (separate histograms for young control and PLX-treated young rats are presented in Fig. S6). The percentage of active PPMPs (of the young rats) deteriorated to a minimum of 50% by the third week post-PPMP implantation (Figs. 8B and S6). Unlike the adult controls, the percentage of active platforms recovered to 100% and the percentage of live channels recovered to 36% on the 6^th^ week post-implantation.

Taken together, the performance of PPMPs implanted to either the control adult rats or adult rats treated by PLX3397 (Figs. 7, 8 and earlier publications from our laboratory [20] and [41]) deteriorated and did not recover spontaneously. In contrast, the recording performance of PPMPs implanted into the young controls or the PLX3397 treated young rats partially recovered after deterioration.

To shed light on the mechanisms underlying these different temporal trends in recording performance, we measured the bulk in-vivo impedances of the system once a week throughout the course of the above described electrophysiological experiments.

To differentiate between the contributions of the electrodes themselves to the measured in-vivo impedance and calculate the changes in net tissue impedance surrounding the implant, we subtracted the net electrode impedances (as measured under in-vitro conditions) from the bulk in-vivo impedances. The in-vivo impedances were measured in anesthetized restrained rats-by passing 1.4 nA through each functional electrode, through the surrounding parenchyma and the bone screw to the ground, using the nanoZ impedance tester (Plexon) at 1 KHz (the pertinent frequency for FP spared). The impedance of the PPMPs themselves were monitored by continuously immersing PPMPs in sterile saline enriched with 10% albumin for 6 weeks using the same parameters as in the in-vivo experiments and a stainless steel bone screw as a counter ground electrode. These measurements revealed that in albumin enriched saline, the microelectrode impedances were maintained at an average value of 1.11±0.12 MΩ (in weeks 1-6,). In contrast, the bulk average in-vivo impedance increased in the three experimental groups during the first two weeks post-implantation by a factor of ~3, from a range of 1±0.05 MΩ on the day of implantation to 3.3± 0.4 MΩ on the second week and further to 3.5 ±0.3 MΩ on the third week post-implantation. Crucially, the average bulk in-vivo impedance in adult rats was significantly (P> 0.001) higher than that of the young rats from the third week and onwards (3.7±0.I5 MΩ, and 2.9±0.4 MΩ respectively). In addition, whereas in the adult rats the bulk impedance remained high for the entire (6 week) recording period, in both the young control and PLX3397-fed young rats, the impedance declined from the third week of implantation (3.2±0.9) and onward to 2.5 ±0.9 MΩ.

These differences in in-vivo impedance kinetics between adult and young cortices correspond temporally to the differences in the density kinetics of the inflammatory microglia as shown above (Figs. 2, 3 and supplementary fig. 2) and to our previous studies on adult rats. Thus, the initial increase in bulk impedance (weeks 1 and 2 post-PPMP implantation) in adult and young rats corresponds in time to the activation, division, migration, adhesion and accumulation of microglia onto the PPMP surface. The decreased impedance in the young control and PLX3397-treated young rats (but not in the adults) corresponds to the disappearance of the microglia adhering to the platform surfaces (Figs. 2 and 3).

## 4. Discussion

This study addressed three related fundamental questions: (1) Can the hypothesis that activated microglia adhering to the surface of implanted PPMPs rather than the multicellular FBR underlie the recording performance deterioration be substantiated? (2) Is the severity of the inflammatory FBR generated by rat cortices age-dependent? (3) Is microglia elimination by CSF1R antagonists more effective in young than adult rats? To address these questions, we examined the temporal relationships between the spatiotemporal densities of microglia, astrocytes and neurons after PPMP implantation, and the recording performance of the implants in young and adult rats under control conditions and after treatment with CSF1R antagonists.

(1) The key shared feature across the recording performance of the implanted control adult rats, the control young rats and PLX3397 treated young rats was the decline in the percentage of the recording platform and live electrodes between the first to the third weeks after implantation (Figs. 7 and 8). Whereas the recording performance of the PPMPs implanted in control adult and PLX5622 treated adult rats as shown in Fig. 7 and [20, 41] remained low for over 6 weeks [see also 20, 41], the performance of the implants in the control young rats and young rats treated by PLX3397 started to recover during the 3^rd^ to 4^th^ week post-PPMP implantation. The initial decline was followed by partial spontaneous recovery of the recording performance in the young rats that occurred within a time window in which the immunohistological observations clearly demonstrated that the astrocyte density, (within the PPMP pores in contact with the implant surface and around it) in shells 1-3 continued to increase (Fig. 3 and S2, and see [20]). Critically recall that in all experimental groups the density of neurons in shells 1-4 (0-100μm from the implant surface) recovered to 75-100% of the control levels and that implanted electrodes can sense FPs generated by neurons residing up to ~150 μm away [21, 61, 62] suggests that the encapsulating astrocytes of adults and young rats did not play a major role in impeding the electrical coupling between the neurons and the electrodes.

In contrast to the astrocytes, the microglia density in both adult and young cortices was high for a period of approximately one week within the platform pores, in contact with it, and in shells 1-7 around it (Figs. 2, 3, S1, S2 and Fig. 8 in [20]). Starting from the second week postimplantation and onward, the microglia density around the implant (NFI and cells/mm^2^) declined spontaneously. The reduced microglia density temporally coincided with the recovery of implant performance. Crucially, the reduction in microglia density differed between the adult and young rats. Whereas in adults, microglia insulation (the PLX resistant subpopulation) remained attached to the implant and within the platform pores for over a >4 weeks after PPMP implantation (Fig. 3, and in [20]), in the control young rats the microglia density (both attached and within the electrodes) dropped to more than 50% as compared to the control adults. In young rats treated by PLX5622 the microglia density was even lower. The persistent presence of microglia attaching to the implanted PPMPs in the control implanted adult rats was temporally correlated with the decline in recording performance of the adult cortical implants (Figs. 6 and 14 in [20]). Importantly, in the PLX5622 treated adult rats in which 95% of the cortical microglia were eliminated, a subpopulation of PLX5622-resistant microglia remained attached to the platform and the recording performance declined just as it did in the control adult rats. Therefore, the PLX5622 chow did not improve the recording performance in adult rats [41]. The recovery of the recording performance in the control young rats and PLX3397-treated young rats and the parallel decrease in the measured bulk impedance were temporally correlated with the reduced densities of the microglia attaching to the PPMPs (Figs. 3, 4, 5, 7 and 8).

Thus overall, the analysis of the temporal relations between the cellular components and the densities of cells comprising the FBR and the recording performance is consistent with the hypothesis that microglia adhering to the implant surface are the major underlying cellular feature that insulates the electrodes from the surrounding cells. Future work is required to generalize the age-related and CSF1R-antagonist effects on the inflammatory cascades induced by PPMP implantations to other types of neuroimplants, by direct experimental testing.

(2) MEA device development and testing for basic brain research, as well as efforts to develop novel bioengineering approaches to mitigate the FBR have been conducted exclusively on adult rodents. Inspired by the TBI literature [46–48], we opted to deviate from the traditional approach and examine whether the FBR severity generated by the cortices of young rats would be ameliorated as compared to adult rats. If borne out, using young rats rather than adult mice and rats for basic research purposes would be advantageous. In addition, understanding the age-dependent molecular and cell biological mechanisms that underlie FBR formation in young and adult rats could contribute to developing ways to curtail the inflammatory cascades.

In contrast to mice, which weigh 20-25 gr in adulthood, young rats weighing ~ 50 gr can be used for electrophysiological brain research since they are large enough to carry remote amplifiers for recording devices that weigh ≤10% of their body weight while behaving freely and demonstrate an advanced behavioral repertoire when only several weeks old (for example, [63, 64]). Since we demonstrated here that the recording performance of PPMPs in young rats is enhanced as compared to adults, comparative molecular and cell biological studies of the mechanisms underlying these differences may help pave the way to mitigating implant-induced inflammation. The vast TBI literature (for review see [65]) documents significant age-dependent differences in the response of microglia to brain insults. These studies have pointed to age-dependent morphological differences, differences in the surveillance and movement of microglia towards injury sites, as well as age-dependent intracellular Ca^2+^ dynamics in response to injury and phagocytic activities. Thus, a comparative study of the FBR to neuroimplants in young and old rat models would be beneficial.

In addition, based on the literature relating BBB breakdown during implantation and FBR formation [66] as well as age-dependent BBB breakdown and recovery it is conceivable that the differences in the rate and effectiveness of the BBB regeneration at the site of probe implantation may show significant age-dependent differences [67, 68]. It is also conceivable that pharmacological approaches that have been tested on adult mice and rats may function significantly better in young rat models and further improve the recording performances of the implants. These include antioxidants that ameliorate CNS oxidative stress and inflammatory processes such as resveratrol [69, 70], melatonin, an antioxidant that protects neurons from excitotoxicity and apoptosis, inhibits pro-inflammatory cytokine production and the activation of NF-?B, and the caspase-1/cytochrome c/caspase-3 cell death pathways [71], as well as dexamethasone, an anti-inflammatory drug [11, 72, 73] and IAXO-101 (Innaxon), an antagonist to the CD14/TLR4 complex, [66, 74].

Finally (3), this study suggests that the CSF1R antagonist provided in young rats’ diets mitigated the FBR and improved the PPMP recording performance in young as compared to adult rats. Future studies, with appropriate control experiments, could apply this finding to basic brain research. Nonetheless, the systemic use of the BBB permeable CSF1R antagonist to improve neuroprobe performance for medical applications is highly curtailed since the elimination of almost the entire microglia population [75] and possibly other cell types is a critical drawback. These limitations can be bypassed in future studies by the use of slow release methodologies or microfluidic applications of the CSF1R antagonist from the implant, which would restrict the elimination of the microglia to the implant vicinity alone.

## 5. Funding

This work was supported by the Israel Science Foundation grant #1808/19. Part of this work was supported by the Charles E. Smith and Prof. Joel Elkes Laboratory for Collaborative Research in Psychobiology.

## 6. Acknowledgments

We thank Plexxikon Inc., for supplying PLX5622. The content is solely the responsibility of the authors and does not necessarily represent the official views of the granting agencies.

## 7. Author Contributions

A.S and M.M.J implanted the platforms. A.S. conducted the electrophysiological recording sessions and analyzed the results. H.E. and A.S prepared and analyzed the immunohistological sections. N.S. remotely coordinated the PPMP fabrication. M.E.S. conceived, designed, and supervised the project, and wrote the manuscript.

## 8. Conflicts of interest

The authors have no conflict of interest related to this research to disclose.

## Notes

### Competing Interest Statement

The authors have declared no competing interest.

