## Supplement Fig. for "Significantly reduced inflammatory foreign-body-response to neuroimplants and improved recording performance in young compared to adult rats"

**17 12 2022**

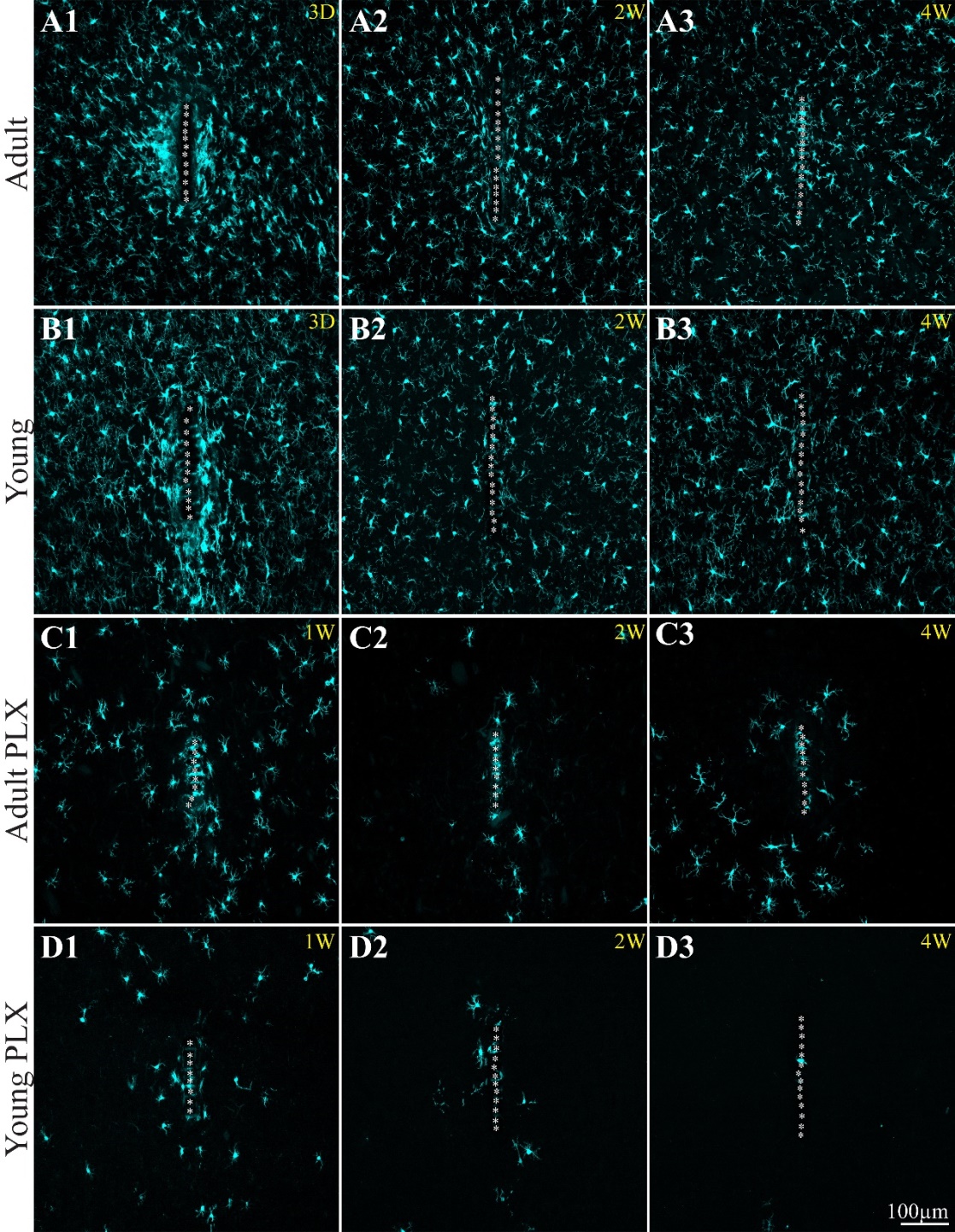
**Supplementary Figures:**

**Fig. S1. Low magnification images comparing the microglia spatiotemporal distribution in contact and around implanted PPMPs in control and PLX5622 treated adult and young rats.** In the shown images the microglia were labeled by the Iba-1 antibody. For the purposes of orientation, the solid polyimide "ridges" in between the pores of the PPMP are labelled by white asterisks. (A1-A3) control adults, on days 3, week 2 and 4 after PPMPs implantations. (B1-B3) control young rats on day 3, weeks 2 and 4 post PPMPs implantations. (C1-C3) PLX5622 treated adults, (D1-D3) PLX5622 treated young rats, 1, 2 and 4 weeks post PPMP implantations. Note that the microglia densities within, in contact and around the implant are larger in the adult (A1-A3) than the young rats (B1-B3, for quantitative information see Figure 3 and Supplementary Figure 2). Ad libitum feeding of the adult and young rats with PLX5622 (starting on the fourth day post PPMP implantation) greatly reduces the microglia densities around the implants. Whereas in the young rats the density of PLX5622-resistant microglia is minimal (D2-D3) in the adults a larger density of PLX5622-resistant microglia remains attached to the platform (C2-C3).

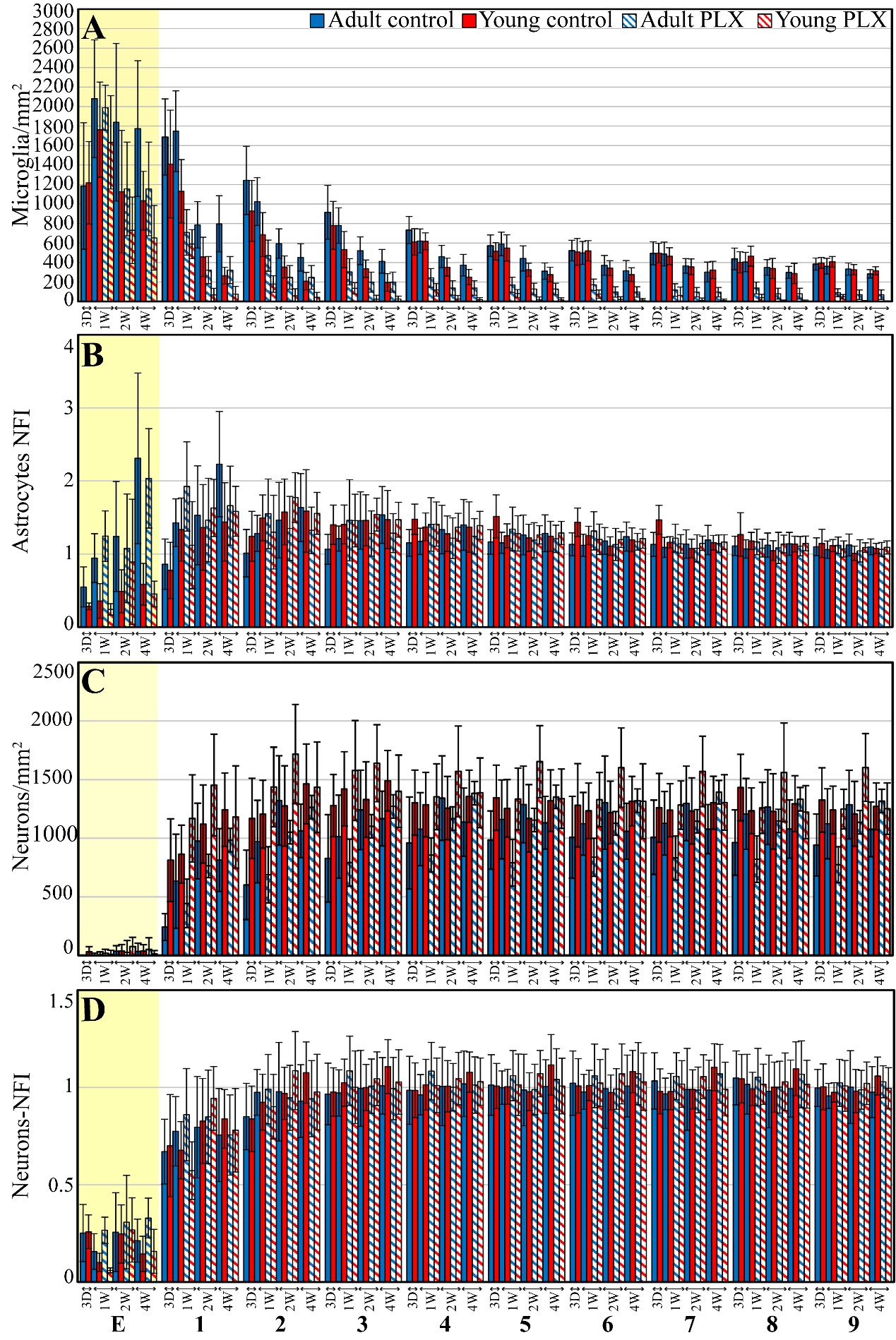

**Fig. S2. The average densities of microglia, astrocytes and neurons as a function of time and over a centripetal distance of 300 µm.** The average densities of the microglia (A), the normalized fluorescent intensity (NFI) of astrocytes (B), neurons density and NFI (C and D, respectively) within the PPMP (yellow background), in contact (shell 1) and around implanted PPMPs (shells 2 to 9) in control and PLX5622 treated adult and young rats, at different points in time post- PPMP implantation. Homogeneous blue columns depict control adult cortices. Homogeneous red columns depict control young rats. Diagonal blue and red stripes depict cortices of adults and young rats fed with PLX5622 4 days after PPMP implantation and thereafter. The times post- platform implantation (3 days, 1, 2, and 4 weeks) and the distance of the measured averages, are given below the X axis of the histograms. Vertical lines correspond to one standard deviation. Black circles indicate significant differences of P<0.01 between the value of control adults and control young rats, triangles indicate significant differences of P<0.01 between PLX5622 treated adults or young rats to control adults or young rats. Similar data depicting the spatiotemporal densities and NFIs values over a centripetal distance of 300µm.

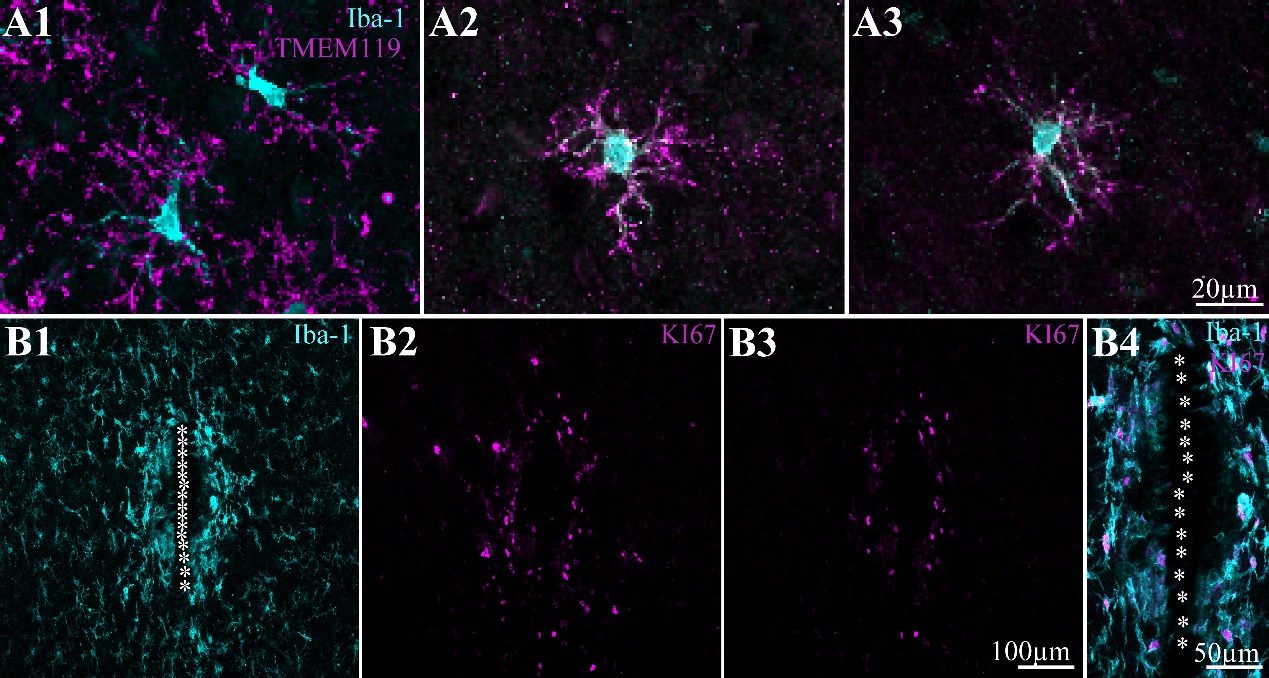
**Fig. S3.** (a) Co-labeling of microglia by Iba-1 (cyan) and TMEM 119 (magenta) in control young rats (A1) and of PLX5622-resistant microglia in young rats fed with PLX5622 for 6 (A2) and 13 days (A3). (B) Microglia mitosis induced by the PPMP implant in young rats. The images were prepared 3 days after implantation. For purposes of orientation, the solid PI “ridges” in between the pores of the PPMP are labeled with white asterisks. (B1) Low magnification of microglia labeled by Iba-1 (cyan). (B2) Mitotic cells residing near the PPMP labeled by KI67 (magenta). (B3) Not all the mitotic cells shown in (B2) are microglia. The shown image was generated after removal of all cells that were not co labeled by Iba-1 and KI67 (B1 and B2). (B4) Enlargement of the merge image of B1 and B3.

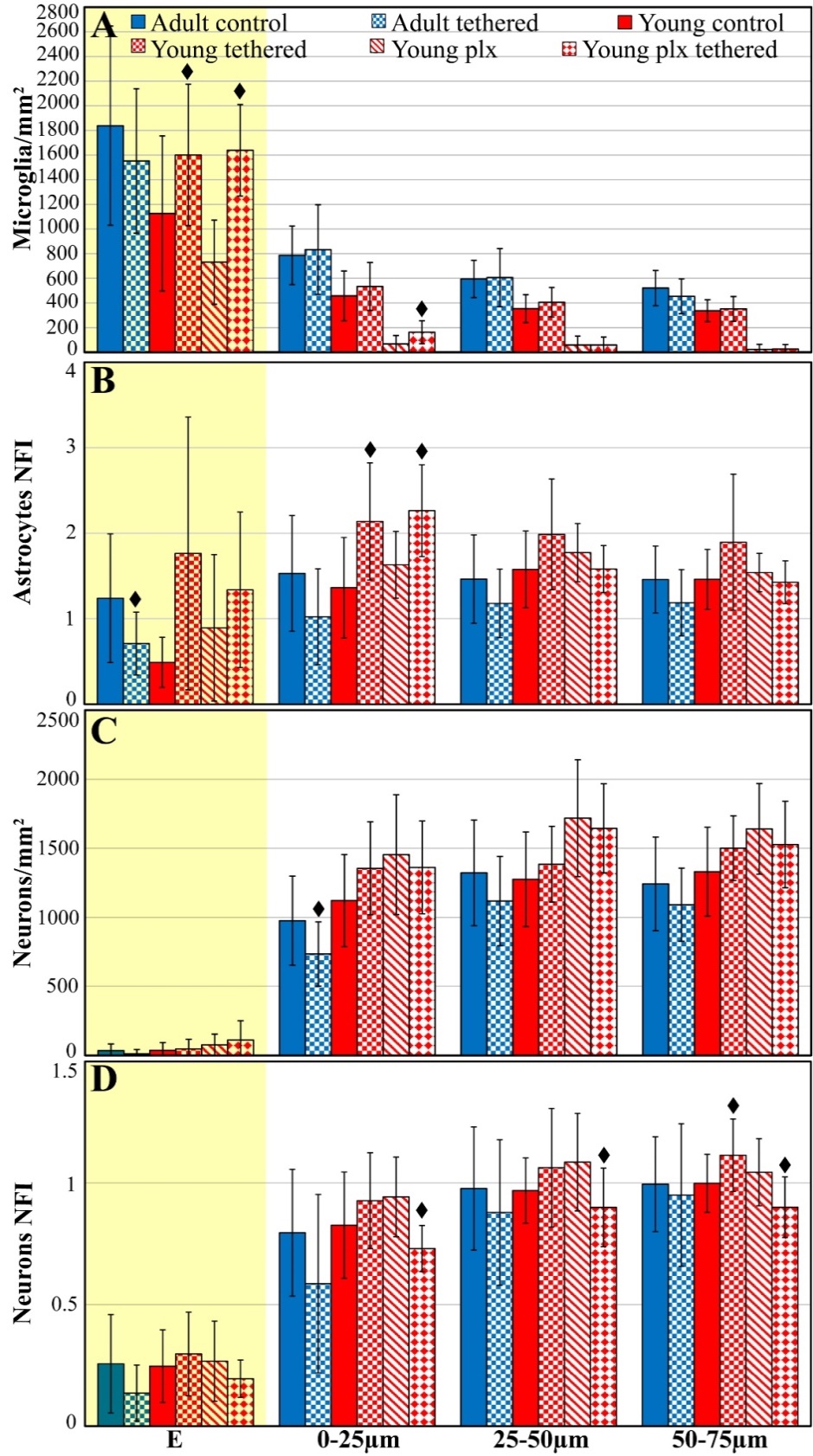

**Fig. S4. The average densities of free floating (untethered) and fixed to the skull (tethered) microglia, astrocytes and neurons, two weeks after implantation within the PPMP and over a centripetal distance of 0-75 µm from the implant surface.** The average densities of the microglia (A), the normalized fluorescent intensity (NFI) of astrocytes (B), neurons density and NFI (C and D, respectively) within the PPMP (yellow background-E), in contact (shell 1) and around implanted PPMPs (shells 2 and 3). Homogeneous blue and red columns depict adult and young control untethered PPMPs. Crossed diagonal blue and red stripes depicts tethered PPMPs in adults and young rats correspondingly. Red diagonals and chattered depicts untethered and tethered PPMPs in young rats fed with PLX5622. Vertical lines correspond to one standard deviation. Black rhombus indicate significant differences of P<0.01 between untethered and tethered PPMPs.

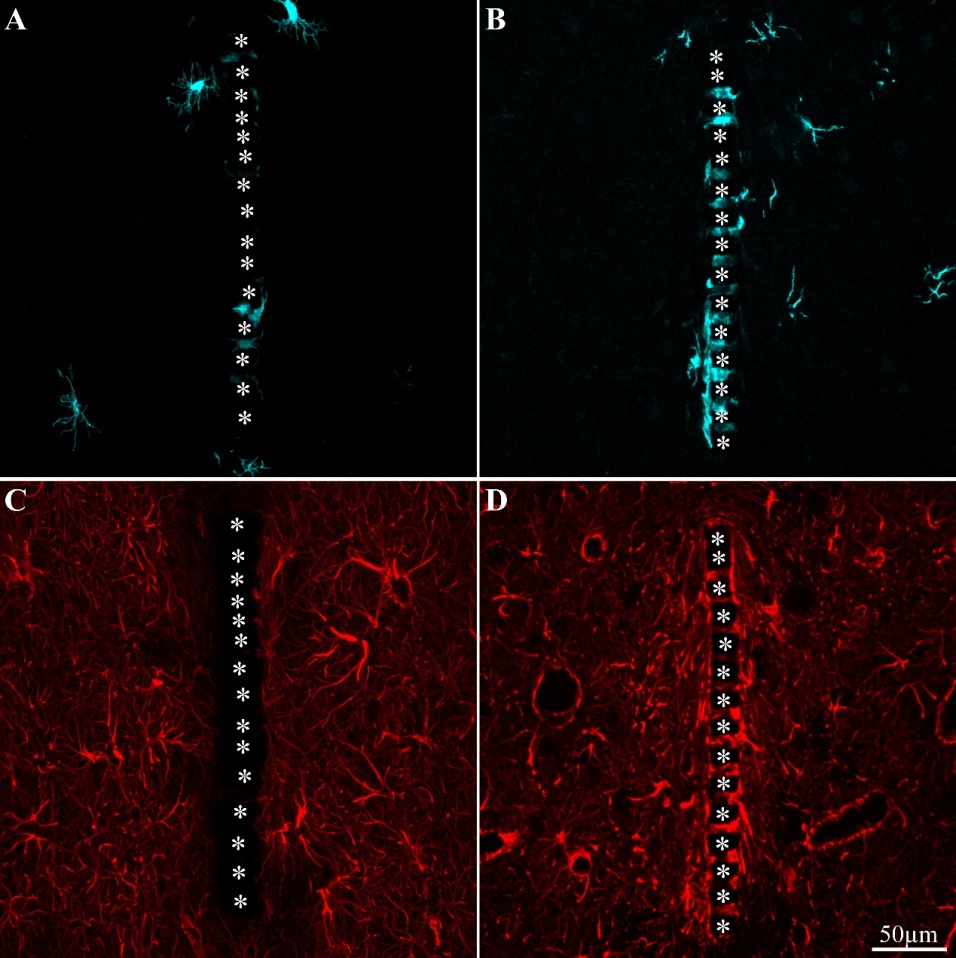

**Fig. S5. Differences in the spatiotemporal distribution of microglia (A, B) and astrocytes (C, D) in contact and around untethered (A and C) and tethered (B and D) PPMPs in young rats fed with PLX5622 two weeks after implantation.** In the shown images the microglia were labeled by the Iba-1 antibody and astrocytes by GFAP. For the purposes of orientation, the solid polyimide "ridges" in between the pores of the PPMP are labelled by white asterisks.

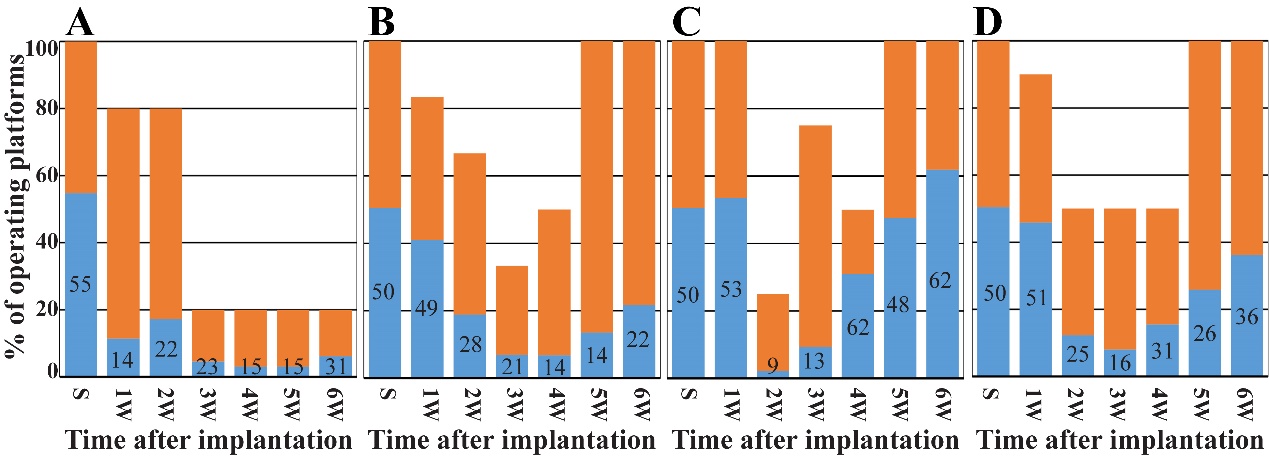
**Fig. S6. Comparison of PPMp performance**. In orange, the percentage of operating platforms, in blue, the percentage of live electrodes. On the day of PPMP implantation 55% of the electrodes were viable in the adult rats (A), and 50% in the young control (B) and young PLX3397 treated rats (C). In the control adult implants (A), the percentage of functional platforms and "live" electrodes decreased with time post implantation. In contrast, in the control young rats (B) and PLX3397 treated young rats (C) after an initial decline of the percentages of active platform and live electrodes the recording performance spontaneously improved. D shows the combined data from all young rats (B&C -control and PLX treated).

**Supplementary Table S1:**

| **Time** | **3 days**  **Adult** | **3 days**  **Young** | **1 week**  **Adult** | **1 week**  **Young** | **2 weeks**  **Adult** | **2 weeks**  **Young** | **4 weeks**  **Adult** | **4 weeks**  **young** |
| --- | --- | --- | --- | --- | --- | --- | --- | --- |
| **µG*** | 25-30/8 | 14/3 | 17/6 | 18/4 | 25/6 | 37/7 | 32/6 | 25/5 |
| **As**# | 23/8 | 6/2 | 13/6 | 17/4 | 39/12 | 36/7 | 31/6 | 22/4 |
| **Ne**# | 24/6 | 14/3 | 16/6 | 21/4 | 34/8 | 37/7 | 27/ 6 | 23/5 |
| **Ne*** | 24/6 | 14/3 | 16/6 | 21/4 | 33/9 | 37/7 | 27/6 | 28/5 |

**Control, untethered**

**PLX 4 days after implantation, untethered**

| **Time** | **2 weeks**  **Adult** | **2 weeks**  **Young** | **2 weeks**  **Young PLX** | **4 weeks**  **Adult** | **4 weeks**  **young** |
| --- | --- | --- | --- | --- | --- |
| **µG*** | 14/4 | 19-24/5 | 18-21/4 | 12/3 | 10-11/2 |
| **As**# | 14/4 | 10-13/5 | 12-15/4 | 12/3 | 6/2 |
| **Ne**# | 14/4 | 17-20/5 | 14-16/4 | 8/3 | 8/2 |
| **Ne*** | 14/4 | 18-23/5 | 17-20/4 | 9/3 | 11/2 |

**Tethered**

| **Time** | **1 week**  **Adult** | **1 week**  **Young** | **2 weeks**  **Adult** | **2 weeks**  **Young** | **4 weeks**  **Adult** | **4 weeks**  **young** |
| --- | --- | --- | --- | --- | --- | --- |
| **µG*** | 16/4 | 22/4 | 34/9 | 20/4 | 22/4 | 25/5 |
| **As**# | 11/4 | 18/4 | 41/10 | 15/4 | 16/4 | 19-21/5-6 |
| **Ne**# | 16/4 | 21/4 | 36/9 | 18/4 | 22/4 | 23/6 |
| **Ne*** | 17/4 | 22/4 | 27-29/8 | 19/4 | 20-22/4 | 23/6 |

Number of examined brain Slices (n) / Hemispheres (N). Ten optical sections were made per a single brain slice. M- Microglia; A- Astrocytes and N- Neurons.

* Cells count, # Normalized Fluorescent Intensity (NFI).

**Supplementary Table S2:**

|  | 3 days Adult | 3 days  Young | 1 week  Adult | 1 week  Young | 2 weeks  Adult | 2 weeks Young | 4 weeks  Adult | 4 weeks  Young |
| --- | --- | --- | --- | --- | --- | --- | --- | --- |
| Microglia* Shells E  Untethered | Control  1185±649 | Control  1218±423 | Control 2080±604  PLX  1989±230 | Control 1763±489  PLX  1633±479 | Control 1839±808  PLX  ▲1155±481  P=0.0006 | Control ●1126±629  P=0.0006  PLX  ▲731±341  P=0.003 | Control 1775±696  PLX  ▲862±450  P=3.3*10^-7^ | Control ●1033±301  P=2.47*10^-6^  PLX  ▲653±332  P=7.04*10^-5^ |
| Microglia* Shells E  Tethered |  |  |  |  | Control 1552±586 | Control ♦1601±574  P=0.007  PLX  ♦1638±371  P=3.47*10^-9^ | Control 1912±605 | Control 893±344 |
| Microglia*  Shells 1 Untethered | Control 1688±391 | Control 1410±553 | Control 1747±415  PLX  ▲709±234  P=2.15*10^-9^ | Control ●1131±326  P=3.35*10^-5^  PLX  ▲589±149  P=1.16*10^-6^ | Control 787±238  PLX  ▲322±139  P=2.07*10^-10^ | Control ●457±202  P=8.83*10^-7^  PLX  ▲68±68  P=2.18*10^-14^ | Control  797±288  PLX  ▲146±82  P=1.32*10^-14^ | Control ●265±87  P=4.64*10^-12^  PLX  ▲75±80  P=9.22*10^-11^ |
| Microglia* Shells 1  Tethered |  |  |  |  | Control 833±364 | Control 534±195  PLX  ♦163±93  P=0.0006 | Control 794±372 | Control ♦496±157  P=0.0005 |
| Astrocytes# Shells E  Untethered | Control  0.55±0.27 | Control ●0.28±0.04  P=0.0001 | Control 0.94±0.33  PLX 1.24±0.35 | Control ●0.36±0.24  P=2.46*10^-5^  PLX 0.2±0.08 | Control 1.24±0.75  PLX 1.08±0.74 | Control ●0.49±0.29  P=4.82*10^-7^  PLX 0.89±0.86 | Control 2.31±1.17  PLX 2.04±0.68 | Control ●0.59±0.29  P=2.82*10^-9^  PLX 0.45±0.18 |
| Astrocytes# Shells E  Tethered |  |  |  |  | Control ♦0.71±0.37  P=0.001 | Control 1.76±1.6  PLX  1.34±0.91 | Control ♦0.85±0.29  P=1.23*10^-7^ | Control 1.18±1.64 |
| Astrocytes# Shells 1  Untethered | Control 0.86±0.34 | Control 0.78±0.38 | Control 1.43±0.33  PLX  1.92±0.61 | Control 1.34±0.43  PLX  1.12±0.6 | Control 1.53±0.68  PLX 1.46±0.57 | Control 1.36±0.59  PLX  1.63±0.39 | Control 2.23±0.72  PLX  ▲1.66±0.54  P=0.004 | Control ●1.44±0.54  P=3.1*10^-5^  PLX  1.58±0.34 |
| Astrocytes# Shells 1  Tethered |  |  |  |  | Control 1.02±0.56 | ♦Control 2.14±0.69  P=0.002  ♦PLX  2.26±0.54  P=0.001 | Control 2.26±1.44 | Control 2.66±0.94 |
| Neurons# Shells E  Untethered | Control 0.25±0.15 | Control 0.26±0.09 | Control 0.16±0.09  PLX  ▲0.27±0.07  P=0.0006 | Control 0.1±0.05  PLX  ▲0.06±0.01  P=0.0008 | Control 0.26±0.2  PLX  0.31±0.24 | Control 0.25±0.15  PLX  0.27±0.16 | Control 0.21±0.11  PLX ▲0.33±0.1  P=0.0002 | Control ●0.14±0.09  P=0.0099  PLX  0.16±0.11 |
| Neurons# Shells E  Tethered |  |  |  |  | Control 0.14±0.12 | Control 0.3±0.17  PLX  0.2±0.08 | Control 0.16±0.07 | Control 0.18±0.11 |
| Neurons# Shells 1  Untethered | Control 0.67±0.17 | Control 0.7±0.26 | Control 0.77±0.18  PLX  0.86±0.24 | Control 0.68±0.15  PLX  0.57±0.15 | Control 0.8±0.26  PLX  0.85±0.24 | Control 0.83±0.22  PLX  0.94±0.16 | Control 0.76±0.24  PLX  0.76±0.21 | Control 0.84±0.15  PLX  0.78±0.21 |
| Neurons# Shells 1  Tethered |  |  |  |  | Control 0.59±0.37 | Control 0.93±0.2  PLX  ♦0.73±0.09  P=6.51*10^-5^ | Control 0.88±0.15 | Control 1±0.25 |
| Neurons* Shells E  Untethered | Control  0±0 | Control  31±44 | Control  4±16  PLX  18±34 | Control  5±25  PLX  10±33 | Control  35±48  PLX  27±59 | Control 38±56  PLX  76±78 | Control 35±70  PLX  51±66 | Control 37±54  PLX  13±29 |
| Neurons* Shells E  Tethered |  |  |  |  | Control  12±31 | Control 47±71  PLX  112±139 | Control  9±26 | Control 46±72 |
| Neurons* Shells 1  Untethered | Control 242±114 | Control ●813±353  P=3.03*10^-5^ | Control 633±401  PLX  4.44±2.08 | Control 864±247  PLX  ▲1169±372  P=0.003 | Control 976±323  PLX  764±361 | Control 1121±333  PLX  ▲1454±434  P=0.007 | Control 814±268  PLX  984±415 | Control ●1243±313  P=1.29*10^-6^  PLX  1181±436 |
| Neurons* Shells 1  Tethered |  |  |  |  | Control  ♦734±233  P=0.007 | Control 1355±337  PLX  1362±335 | Control  896±258 | Control 1083±186 |

Mean ± one standard derivation of the average number of cells per mm^2^ (* microglia and neurons) or Normalized Fluorescent Intensity (NFI, # for astrocytes and neuron cell bodies and neurites) within the implanted platform (referred to as Shell E) and 0-25µm away from the platform’s surface (Shell 1). T-test was conducted for two- samples assuming unequal variances. ●, ▲, ♦: P<0.01. ● Adults versus young rats; ▲ Control versus PLX5622 fed rats; ♦ Tethered versus untethered implants.
